# Small-field visual projection neurons detect translational optic flow and support walking control

**DOI:** 10.1101/2023.06.21.546024

**Authors:** Mathew D. Isaacson, Jessica L. M. Eliason, Aljoscha Nern, Edward M. Rogers, Gus K. Lott, Tanya Tabachnik, William J. Rowell, Austin W. Edwards, Wyatt L. Korff, Gerald M. Rubin, Kristin Branson, Michael B. Reiser

## Abstract

Animals rely on visual motion for navigating the world, and research in flies has clarified how neural circuits extract information from moving visual scenes. However, the major pathways connecting these patterns of optic flow to behavior remain poorly understood. Using a high-throughput quantitative assay of visually guided behaviors and genetic neuronal silencing, we discovered a region in *Drosophila*’s protocerebrum critical for visual motion following. We used neuronal silencing, calcium imaging, and optogenetics to identify a single cell type, LPC1, that innervates this region, detects translational optic flow, and plays a key role in regulating forward walking. Moreover, the population of LPC1s can estimate the travelling direction, such as when gaze direction diverges from body heading. By linking specific cell types and their visual computations to specific behaviors, our findings establish a foundation for understanding how the nervous system uses vision to guide navigation.

## INTRODUCTION

The global pattern of perceived visual motion, termed optic flow^1^, provides rich information about the structure of an observer’s local environment. The fruit fly, *Drosophila melanogaster*, is a prominent model system for establishing the circuit implementation of visual computations and for linking these computations to behavioral control^2^. When flies move through an environment, the pattern of optic flow they experience contains information about their own movement, the layout of the surrounding environment, and the relative movement of nearby objects in their field of view^1, 3^. These motion cues are used to stabilize the fly’s heading^4, 5^, track objects^6, 7^, or avoid collisions^8, 9^, and these different behavioral reactions have long suggested that distinct neuronal circuits can distinguish between different patterns of motion.

Much progress has been made towards identifying the circuits and algorithms that compute the local direction of motion in *Drosophila*^10–15^. In the fly visual system, two 4^th^ order neurons, T4 and T5, are the first cells that exhibit directionally selective signals. T4 neurons encode the motion of bright (ON) edges, while T5 encode the motion of dark (OFF) edges. The four subtypes of T4 and T5 neurons are each selective for one cardinal direction and terminate in one of four retinotopic layers of the lobula plate^16–19^. Several groups have used genetic targeting of individual cell types in the visual motion pathway and measured robust, specific effects of neuronal silencing on visual motion guided behaviors^6, 20, 21^. However, these studies have generally analyzed the contributions of early visual system neurons up to and including T4 and T5. Establishing causal links between visual processing by output neurons of the lobula plate or their targets in the central brain, and behavioral control has proven far more challenging, likely because these outputs are a large, diverse set of neurons, most of which cannot be routinely accessed using precise genetic tools and have therefore not been carefully examined.

A group of neurons spanning large regions of the lobula plate, the Lobula Plate Tangential (LPT) neurons, have been extensively characterized in larger flies, most notably in *Calliphora*. The lobula plate is organized retinotopically, so neurons with large dendritic arbors are expected to respond to visual motion over large parts of the eye. Indeed, this group of uniquely identifiable neurons encode ‘wide field’ optic flow and project to the central brain^22, 23^. The best characterized LPTs, the HS (horizontal system) and VS (vertical system) neurons, are primarily tuned to the pattern of visual motion induced by rotations of the body and/or head^24^. The HS and VS cells have also been characterized in *Drosophila*^25^. However, silencing these rotation-selective neurons using genetic methods has not eliminated the behavioral turning responses to rotational visual motion^26^, but has revealed effects on head movements^27^. One possibility is that these large neurons are specialized for gaze-stabilizing head control movements^27–29^, consistent with their tuning for rotational movement, while groups of cells distinct from the well-studied VS and HS cells transduce the visual signals used for body navigation.

While rotational optic flow is clearly important for stabilizing an animal’s gaze and heading direction, detecting and regulating translational optic flow is critical for navigating through an environment, as occurs in goal-directed behavior towards objects such as conspecifics, food sources, or refuge^1, 3^. Yet very little is known about neurons specialized for encoding the flow fields arising from translational motion. Such neurons are presumably essential for regulating forward locomotion, an exploratory behavior whose visual control is surprisingly complex^5, 9, 30–32^. Using genetic silencing, the visual control of forward-walking has been attributed to the T4/T5 pathway^33^, but these neurons only detect motion within a small field of view, so it is not known which neurons downstream of T4/T5 are selective for translational motion and participate in regulating forward locomotion. In blowflies, several cells (presumed to be LPTs) have been recently discovered to be selective for the visual motion corresponding to sideways and upwards translation^34^. However, no forward or backward translation-selective neurons have been described. In addition to the large LPTs, small field neurons projecting from the lobula plate to ‘optic glomeruli’ in the central brain have been described in blowflies^35^ (and recently found in *Drosophila* EM connectome data^36^), but due to their small size, the analysis of their functional properties has not been possible in large flies.

In this study, we describe a systematic approach to mapping neurons required for the visual regulation of locomotion in *Drosophila melanogaster*. We first set out to identify candidate brain regions required for the visual control of walking with a neuroanatomical silencing screen making use of a high-throughput assay. Our approach is analogous to the loss-of-function genetic screens that have been used extensively in *Drosophila* to identify genes required for particular functions^37^. Applying a recently established analytical method we found a specific region in the protocerebrum as a critical step in the transduction of visual signals into behavioral actions. We then identified several small-field Visual Projection Neurons (VPNs) innervating this region and developed specific genetic tools for targeting these cells. From this detailed anatomical analysis, we identified a single cell type, LPC1, that we further investigated using neuronal silencing, activation, and calcium imaging. This cell type strongly links the detection of translational optic flow to the control of forward walking, providing a foundation for understanding how visual motion is processed to regulate behavior.

## RESULTS

### Localizing the neuroanatomical correlates of visuomotor behavior

To identify neurons that contribute to transforming visual motion to behavior, we developed a platform that integrates multiple visuo-motor behaviors, individually established in previous studies^32, 38–40^, that was optimized for high-throughput operation. In our Fly Vision Box, groups of 10-15 flies were placed in clear plastic corridors (tubes) inside of a temperature-controlled, light-tight box. The box holds 6 such corridors. Along either side of each tube, a row of 64 green LEDs presented drifting visual motion patterns. Each tube was capped with a transparent block such that the flies could see a pair of green and UV LEDs, aligned with the ends of the tubes, that were illuminated to examine phototaxis and color preference behavior. The box also contained a pair of vibrating motors that were used to startle the flies at the start of each trial. An on-board microcontroller scheduled events, at 1 ms intervals, pre-specified in an experimental protocol— including the activation of different LEDs and triggers for each camera frame. An infrared (IR) backlight below the corridors enabled high-contrast visualization of the flies walking in the transparent tubes. Videos were acquired at 25 Hz with an IR-sensitive camera mounted above, that recorded through an IR-pass window that blocked visible light from entering (further details in methods; Fig. 1A). Offline tracking of the flies in the recorded videos was used to measure the reactions of the group of flies within each tube. The summarized walking behavior from a typical experiment shows the consistent directional reactions of the flies to gratings moving with different speeds and in opposite directions (Fig. 1B). In developing the assay, we were impressed by two features of the visual motion response (Fig. 1C, S1A): (1) for a given genotype, the averaged behavior of groups of flies was remarkably consistent across experiments, and (2) the flies’ behavior is tuned to the speed of the visual grating pattern with a temporal frequency tuning—peaking between 8 and 20 Hz—that agrees well with measurement from much more laborious experiments, such as single-fly tethered flight^41, 42^ or walking^13, 43^. A further, noteworthy feature of these responses is that flies exhibited a ‘reverse-optomotor’ response, that is they preferred to walk not with, but against the direction of visual motion. Such a response has been described in similar assays^32^ but is at odds with predictions based on the so-called (syn-directional) ‘optomotor’ behavior that is routinely studied in response to rotational visual motion^4^.

**Fig. 1.**
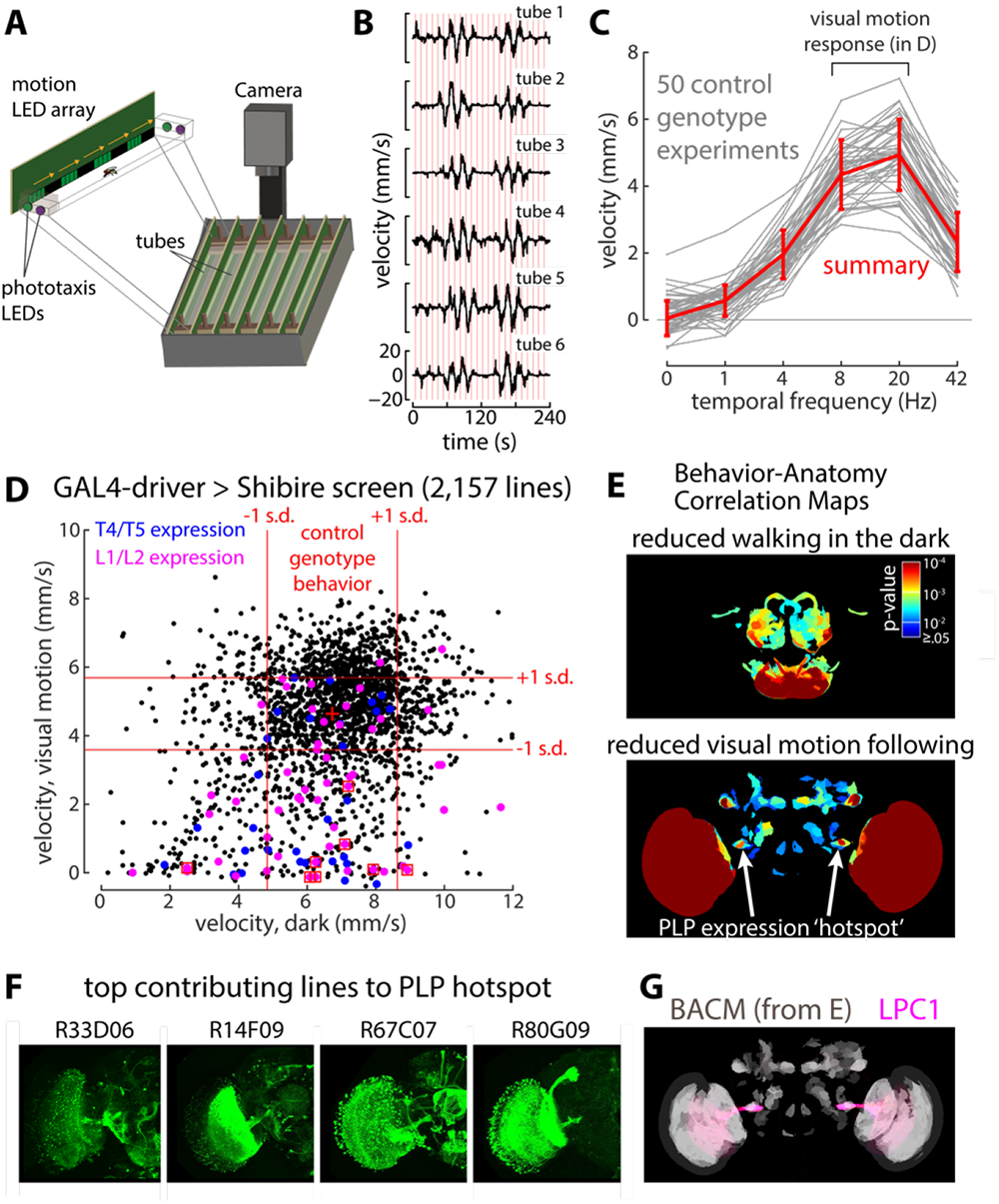
Localizing the neuroanatomical correlates of visuomotor behavior. (A) Left: Schematic of the Fly Vision Box assay. 10-15 flies are placed in each of 6 small clear plastic corridors and presented with motion stimuli by LEDs lining one of the walls of each corridor. The direction of scrolling motion causes a directional walking response, measured by infrared cameras mounted above and off-line fly movement tracking and analysis. (B) Representative experiment of walking flies, of the control genotype (‘empty’ GAL4 line crossed to *UAS-Shibire^ts^*^1^) used as a reference for the GAL4 line screen, in the Fly Vision Box. The timeseries of walking responses to drifting gratings presented at multiple speeds (start of each trial indicated by vertical red lines), plotted as the mean velocity of all flies in each tube (details of these ∼4-minute protocols in the Methods). These timeseries are summarized as tuning curves across the temporal frequency of the grating motion in each trial, in (C), which shows the aggregate responses of control flies run in 50 independent experiments, selected at random, to demonstrate the consistency of these behavioral reactions. The N = 729 control experiments are further summarized (in red) as the mean ± s.d., which is used as a basis for comparing the effects of silencing neurons in the various GAL4 lines (see also Fig. S1A). 2157 ‘generation 1’ GAL4 lines (4144 experiments, nearly all genotypes run twice) driving expression of *UAS-Shibire^ts^*^1^ were screened through the Fly Vision Box. The results of this screen are summarized (D), where each point is the mean of the behavioral metric across all tubes and all experiments for one genotype. This scatter plot shows the velocity of flies following the visual motion of the drifting grating (mean response to 8 and 20 Hz conditions, as indicated in C) against the mean walking velocity in the dark before any visual motion was presented (to capture general deficits in locomotion). Red lines mark ±1 s.d. from the mean of these metrics for the control genotype (mean indicated by red cross). Lines with confirmed expression in the Lamina Monopolar Cells L1 or L2 are indicated in magenta (red boxes indicate expression in both) and lines with T4/T5 expression are indicated in blue. (E) The brain regions of GAL4-diver expression that are correlated with specific behavioral differences, relative to control flies, represented as maximum intensity projections of Behavior Anatomy Correlation Maps (BACMs)^46^. BACMs for reduced walking in the dark (top) and following of fast visual motion (bottom) show very different units of driver-line expression. The BACMs are color coded by p-values (on a log plot, see Methods). Arrows point to the anatomical region that will be the focus of much of this manuscript, a ‘hotspot’ of expression in the Posterior Lateral Protocerebrum (PLP). Related BACMs are shown in Fig. S1D-F. (F) Representative images of expression patterns (where the indicated GAL4 line drives *UAS-GFP*, shows as maximum intensity projections) for four lines that strongly contribute to the hotspot identified in (E). (G) The hotspot, seen in the BACM from (E), shown in gray, strongly overlaps with a cell type called the Lobula Plate Columnar type 1 cell, represented here by the expression pattern of a split-GAL4 line (shown in magenta, detailed in Table 1) developed for this study. The genotypes used in the GAL4 line screen are in Table S1.

**Table 1.** List of genotypes (outside of the GAL4 screen) used throughout the manuscript.

| Genotype details for all anatomy figures |  |  |  |
| --- | --- | --- | --- |
| Figure | Driver line(s) and cell type targeted | Reporter transgene(s) | Description |
| 1F | *R33D06,R14F09,R67C07,R80G09* | *pJFRC2-10XUAS-IVS-mCDGFP* | GAL4 line images from Jenett et al |
| 1G | *SS02575 (R81A05-p65AD (attP40); VT031495-DBD (attP2))* (LPC1) | *pJFRC51-3XUAS-IVS-syt::smHA (su(Hw)attP1)*, *pJFRC225-5XUAS-IVS-myr::smFLAG (VK00005)* (only myr::smFLAG pattern shown) | Overlay of LPC1 cells with BACM |
| 2A | See 2,B,C,D,E | See 2B,C,D,E | Composite of images in 2B,C,D,E with reference brain pattern (63x objective) |
| 2B | *SS02575 (R81A05-p65AD (attP40); VT031495-DBD (attP2))* (LPC1) | *pJFRC51-3XUAS-IVS-syt::smHA (su(Hw)attP1)*, *pJFRC225-5XUAS-IVS-myr::smFLAG (VK00005)* (only myr::smFLAG pattern shown) | Composite of registered images with reference brain pattern (63x objective) |
| 2C | *SS25868 (VT032900-p65AD (attP40); VT016114-DBD (attP2))* (LPC2) | *pJFRC51-3XUAS-IVS-syt::smHA (su(Hw)attP1)*, *pJFRC225-5XUAS-IVS-myr::smFLAG (VK00005)* (only myr::smFLAG pattern shown) | Composite of registered images with reference brain pattern (63x objective) |
| 2D | *SS02407 (VT057342-p65AD (attP40); VT044492-DBD (attP2))* [LLPC1, (LLPC2)] | *pJFRC51-3XUAS-IVS-syt::smHA (su(Hw)attP1)*, *pJFRC225-5XUAS-IVS-myr::smFLAG (VK00005)* (only myr::smFLAG pattern shown) | Composite of registered images with reference brain pattern (63x objective) |
| 2E | *SS02436 (VT029598-p65AD (attP40); VT046081-DBD (attP2))* (LLPC2,LLPC3) | *pJFRC51-3XUAS-IVS-syt::smHA (su(Hw)attP1)*, *pJFRC225-5XUAS-IVS-myr::smFLAG (VK00005)* (only myr::smFLAG pattern shown) | Composite of registered images with reference brain pattern (63x objective) |
| 2F | *SS02575* (LPC1) | MCFO-1 [*pBPhsFlp2::PEST (attP3)*];; *pJFRC201-10XUAS-FRT>STOP>FRT-myr::smGFP-HA (VK00005)*, *pJFRC240-10XUAS-FRT>STOP>FRT-myr::smGFP-V5-THS-10XUASFRT>STOP>FRT-myr::smGFP-FLAG (su(Hw)attP1)*] | Composite of registered images with reference brain pattern; cells from multiple specimens (63x objective) |
| 2G | *SS25868* (LPC2), *SS02436* (LLPC2,LLPC3), *SS02437 (VT048351-p65AD (attP40); VT057342-DBD (attP2))* [LLPC1,(LLPC2)], *SS02575* (LPC1) | *pJFRC51-3XUAS-IVS-syt::smHA (su(Hw)attP1)*, *pJFRC225-5XUAS-IVS-myr::smFLAG (VK00005)* (only myr::smFLAG pattern shown) | Single slices of composite image stack of registered images and reference brain; different channel combinations shown (63x objective) |
| 2H | *SS00324 (R59E08-p65AD (attP40); R42F06-DBD (attP2))* (T4/T5), *SS02575* (LPC1) | *pJFRC51-3XUAS-IVS-syt::smHA (su(Hw)attP1)*, *pJFRC225-5XUAS-IVS-myr::smFLAG (VK00005)* (*SS00324*, only syt::smHA pattern shown), MCFO-1 (*SS02575*) | Composite of registered images with reference brain pattern (63x objective) |
| 2I | *SS02575* (LPC1), *SS25868* (LPC2), *SS02406* (LLPC1), *SS02436* (LLPC2,LLPC3) | MCFO-1 | MCFO-labeled cells with anti-Brp (mAb Nc82) labeling<br>Projections through resampled, cropped volumes of the original confocal stacks (63x objective) |
| 2J | SS02575 (LPC1) | MCFO-1 | Composite of registered images with reference brain pattern; cells from multiple specimens (63x objective) |
| S2A | SS25868 (LPC2) | MCFO-1 | Composite of registered images with reference brain pattern (63x objective) |
| S2B | SS02437(LLPC1) | MCFO-1 | Composite of registered images with reference brain pattern (63x objective) |
| S2C | SS02422 (*VT046081-p65AD (attP40); VT029598-DBD(attP2)*) (LLPC2) | MCFO-1 | Composite of registered images with reference brain pattern (63x objective) |
| S2D | SS02422 (LLPC3) | MCFO-1 | Composite of registered images with reference brain pattern (63x objective) |
| S2E | SS25868 (LPC2) | MCFO-1 | Composite of registered images with reference brain pattern; cells from multiple specimens (63x objective) |
| S2F | SS02406 (LLPC1) | MCFO-1 | Composite of registered images with reference brain pattern; cells from multiple specimens (63x objective) |
| S2G | SS02436 (LLPC2), SS02422 (LLPC2) | MCFO-1 | Composite of registered images with reference brain pattern; cells from multiple specimens (63x objective) |
| S2H | SS02436 (LLPC3) , SS02422 (LLPC3) | MCFO-1 | Composite of registered images with reference brain pattern; cells from multiple specimens (63x objective) |
| 3B,5A,6 A | SS02575 (LPC1), SS025868 (LPC2), SS02407 [LLPC1,(LLPC2)], SS02437 [LLPC1,(LLPC2)], SS22436 (LLPC2,LLPC3) | *pJFRC51-3XUAS-IVS-syt::smHA (su(Hw)attP1)* ,*pJFRC225-5XUAS-IVS-myr::smFLAG (VK00005)* (only myr::smFLAG pattern shown) | Maximum intensity projection with anti-Brp (mAb Nc82) labeling (20x)<br>Figure 5A/6A include SS02575/SS02407 images from Figure 3B |
| S3B | LPC1 drivers: SS02700 (R81A05-p65AD (attP40); VT043014-DBD(attP2)), SS00377(R81A05-p65AD (attP40); R72G06-DBD(attP2)) | *pJFRC51-3XUAS-IVS-syt::smHA (su(Hw)attP1)* ,*pJFRC225-5XUAS-IVS-myr::smFLAG (VK00005)* (only myr::smFLAG pattern shown) | Maximum intensity projection with anti-Brp (mAb Nc82) labeling (20x objective) |
| S3C | SS02575, SS25886, SS02407, SS22437, SS22436, , SS00377 | *pJFRC51-3XUAS-IVS-syt::smHA (su(Hw)attP1)* ,*pJFRC225-5XUAS-IVS-myr::smFLAG (VK00005)* (only myr::smFLAG pattern shown) | Maximum intensity projection with anti-Brp (mAb Nc82) labeling (20x objective) |
| 4D | SS02575 (LPC1) | *w<sup>+</sup>,pJFRC300-20XUAS-FRT>-dSTOP-FRT>-CsChrimson-* | Maximum intensity projection with anti-Brp (mAb Nc82) labeling |
|  |  | *mVenus(attP18),pBPhsFlp2::PEST(attP3)* | (20x objective) |
| 7A | *SS02575* (LPC1), *SS02407* (LLPC1,(LLPC2)), *SS34215* (*VT002033* -*p65AD* (*attP40*); *R78A06-DBD* (*attP2*)) (LPC-IN) | *pJFRC51-3XUAS-IVS-syt::smHA* (*su(Hw)attP1*), *pJFRC225-5XUAS-IVS-myr::smFLAG* (*VK00005*) (only myr::smFLAG pattern shown for *SS34215*; illustrated glomeruli are based on syt::smHA patterns of *SS2575* and *SS02407*) | Composite of registered images with reference brain pattern and glomerulus labels<br>(63x objective) |
| 7B | *SS34215* (LPC-IN) | *pJFRC51-3XUAS-IVS-syt::smHA* (*su(Hw)attP1*), *pJFRC225-5XUAS-IVS-myr::smFLAG* (*VK00005*) | Composite of registered images with reference brain pattern<br>(63x objective) |
| 7C,C' | See 7B | See 7B | Single confocal slices (63x objective) |
| 7D,D' | *SS34215* (LPC-IN), *MI09277-LexA:QFAD* (GAD1-LexA) | *pJFRC19-13XLexAop2-IVS-myr::GFP* (*su(hw)attP8*), *pJFRC21-10XUAS-IVS-mCD8::RFP* (*attP18*) | Single confocal slices (63x objective) |
| S8 | *SS034215* | *UAS-7xHaloTag::CAAX* (*VK00005*)<br>GAD1, ChAT, VGlut FISH probes | Single confocal slices (63x objective) |
| **Genotypes for the behavior and Calcium imaging experiments** |  |  |  |
| 3A, 4B-C, S4 | *w; BPP65ADZp* (*attP40*); *BPZpGdbd* (*attP2*)<br>(enhancerless split GAL4) | *w+(DL);;UAS-Shibire<sup>ts1</sup>* | Shibire (silencing) control genotype |
| 3A, 4B-C, S4 | *SS02575* (LPC1) | *w+(DL);;UAS-Shibire<sup>ts1</sup>* | Shibire (silencing) |
| 3A | *SS25868* (LPC2) | *w+(DL);;UAS-Shibire<sup>ts1</sup>* | Shibire (silencing) |
| 3A,S6 | *SS02407* (LLPC1, some LLPC2) | *w+(DL);;UAS-Shibire<sup>ts1</sup>* | Shibire (silencing) |
| 3A | *SS02437* (LLPC1, LLPC2) | *w+(DL);;UAS-Shibire<sup>ts1</sup>* | Shibire (silencing) |
| 3A | *SS02436* (LLPC2, LLPC3) | *w+(DL);;UAS-Shibire<sup>ts1</sup>* | Shibire (silencing) |
| S3A | *w; BPP65ADZp* (*attP40*); *BPZpGdbd* (*attP2*)<br>(enhancerless split GAL4) | *w+(DL);UAS-Kir2.1;UAS-Shibire<sup>ts1</sup>* | Shibire + Kir (double-silencing) control genotype |
| S3A | *SS02575* (LPC1) | *w+(DL);UAS-Kir2.1;UAS-Shibire<sup>ts1</sup>* | Shibire + Kir (double-silencing) |
| S3A | *SS00377* (LPC1) | *w+(DL);UAS-Kir2.1;UAS-Shibire<sup>ts1</sup>* | Shibire + Kir (double-silencing) |
| S3A | *SS02700* (LPC1) | *w+(DL);UAS-Kir2.1;UAS-Shibire<sup>ts1</sup>* | Shibire + Kir (double-silencing) |
| 4D | *SS02575* (LPC1) | *w+,pJFRC300-20XUAS-FRT>-dSTOP-FRT>-CsChrimson-mVenus(attP18),pBPhsFlp2::PES T* (*attP3*) | Chrimson (Unilateral and bilateral activation and control) |
| 5B-E<br>S5C,D | *SS02575* (LPC1) | *w+(DL); pJFRC7-20XUAS-IVS-GCaMP6m* (*VK00005*) | GCaMP6m (imaging) |
| 6B-E,<br>S7 | *SS02407* (LLPC1, some LLPC2) | *w+(DL);; pJFRC7-20XUAS-IVS-GCaMP6m* (*VK00005*) | GCaMP6m(imaging) |
| 7E | *SS34215* (LPC1-IN) | *w+(DL);; 20XUAS-Syn-OpGCaMP6s* (*su(Hw)attP1*) | OpGCaMP6s(imaging) |

2,157 ‘generation 1’ GAL4 driver lines^44^, were crossed to *UAS-Shibire^ts^*^1^, a temperature-sensitive ortholog of dynamin that blocks chemical synaptic transmission at elevated temperatures^45^, and the F1 offspring were successfully run through a compact protocol in the Fly Vision Box (warmed to 34° C; nearly all lines were run multiple times, 4144 total experiments). Many lines exhibited reduced visual and/or walking responses when compared to the control genotype (an ‘empty’ GAL4 line crossed to Shibire^ts1^ run throughout the screen for a total of 729 experiments; Fig. 1D). For example, 540 lines showed a reduced response to visual motion (below the lower horizontal red line; < mean – 1 s.d. of the control response). The set of lines screened were selected from a larger collection of transgenic lines whose expression patterns were previously characterized^44^. While the lines drive expression in neurons throughout the brain, for some lines we could readily identify expression in specific visual neuron types, for example in L1 and L2, the Lamina Monopolar Cells that convey photoreceptor inputs into the motion pathway, and in T4/T5, the directionally selective cells. As expected, most (>2/3) of the lines we identify with expression in these cells exhibited visual motion deficits, and stronger expression, such as in both L1 and L2 together, lead to even stronger phenotypes (Fig. 1D). However, this approach to analyzing these results is limited since (1) not all lines expressing in a particular cell type produced consistent phenotypes, (2) many lines without expression in “obvious” candidate cell types showed behavioral deficits, and (3) using current methods, the identification of all cell types expressed in 1000s of GAL4 lines is a nearly impossible task.

To overcome these limitations, we used a recently developed analytical method^46^, that identifies anatomical components of expression patterns that correlate with specific behavioral phenotypes. In the previous study, this analysis was developed for correlating locomotion and social behavior phenotypes to patterns of activated neurons^46^, while here we apply the identical experimental framework (using the software developed in the previous study and a nearly overlapping set of GAL4 driver lines) for correlating visual behavior phenotypes to expression patterns of neurons with suppressed synaptic transmission. The outcome of this analysis is a Behavior-Anatomy Correlation Map (BACM) for each behavioral metric. Two such maps, one for reduction in walking speed in the dark, and a second for the directional response to fast motion, are shown in Fig. 1E (with related BACMs in Fig. S1E-G, see methods for details of BACM generation). As was seen in many BACMs from the previous study^46^, different behavioral metrics have very different corresponding maps. Reduced walking in the dark correlates with silencing neurons with expression in the Suboesophageal Zone (SEZ), central complex, and the antenna lobes. As expected, the BACM for reduced visual motion responses shows a strong contribution from optic lobe expression, as well as multiple ‘hot spots’ in the central brain. Among these, we identified a subregion of the posterior lateral protocerebrum (PLP). While the labeling of this PLP region in the BACM could represent the aggregate contribution of many cell types, we nevertheless wondered whether a single pathway, or possibly a single cell type, might disproportionately account for the behavioral deficits associated with this structure. This ‘hotspot’ (white arrows in Fig. 1E) is distinct from the optic glomeruli receiving input from columnar lobula VPNs^47–50^ and also did not match the location of the terminals of HS and VS axon terminals, suggesting contributions of other cell types. We examined the expression patterns of GAL4 lines that contributed to this this feature of the BACM (some examples in Fig. 1F) and found that several included populations of large numbers of similar neurons projecting from the lobula plate to the PLP (one such population shown with overlap, Fig. 1G). Small field lobula plate VPNs have been reported in multiple fly species^16, 35, 36, 47, 48^, including *Drosophila*, but their function and detailed anatomy is only recently being explored^9, 18, 36, 51^.

### Small-field visual projection neurons of the Lobula Plate

As a basis for the further study of small-field lobula plate projection neurons and their possible contributions to visual motion guided behavior, we first examined potential subtypes of this group. Using anatomical criteria, we were able to distinguish five types of columnar lobula plate projection neurons (Fig. 2), considerably more than previously described by light microscopy in *Drosophila*^47, 48^. This analysis preceded the recent completion and cataloging of cell types in the Hemibrain EM volume^36^,which corroborated the presence and general projection pattern of the five cell types in the central brain. Subsequently, a detailed study of lobula plate connectivity further corroborated the presence of 5 distinct small-field cell-types based on innervation patterns of that neuropil^18^. Our light microscopy analysis was important for the annotation of these neurons in both EM volumes, which only contain a small part of the optic lobe or lack the central brain, respectively.

**Fig. 2.**
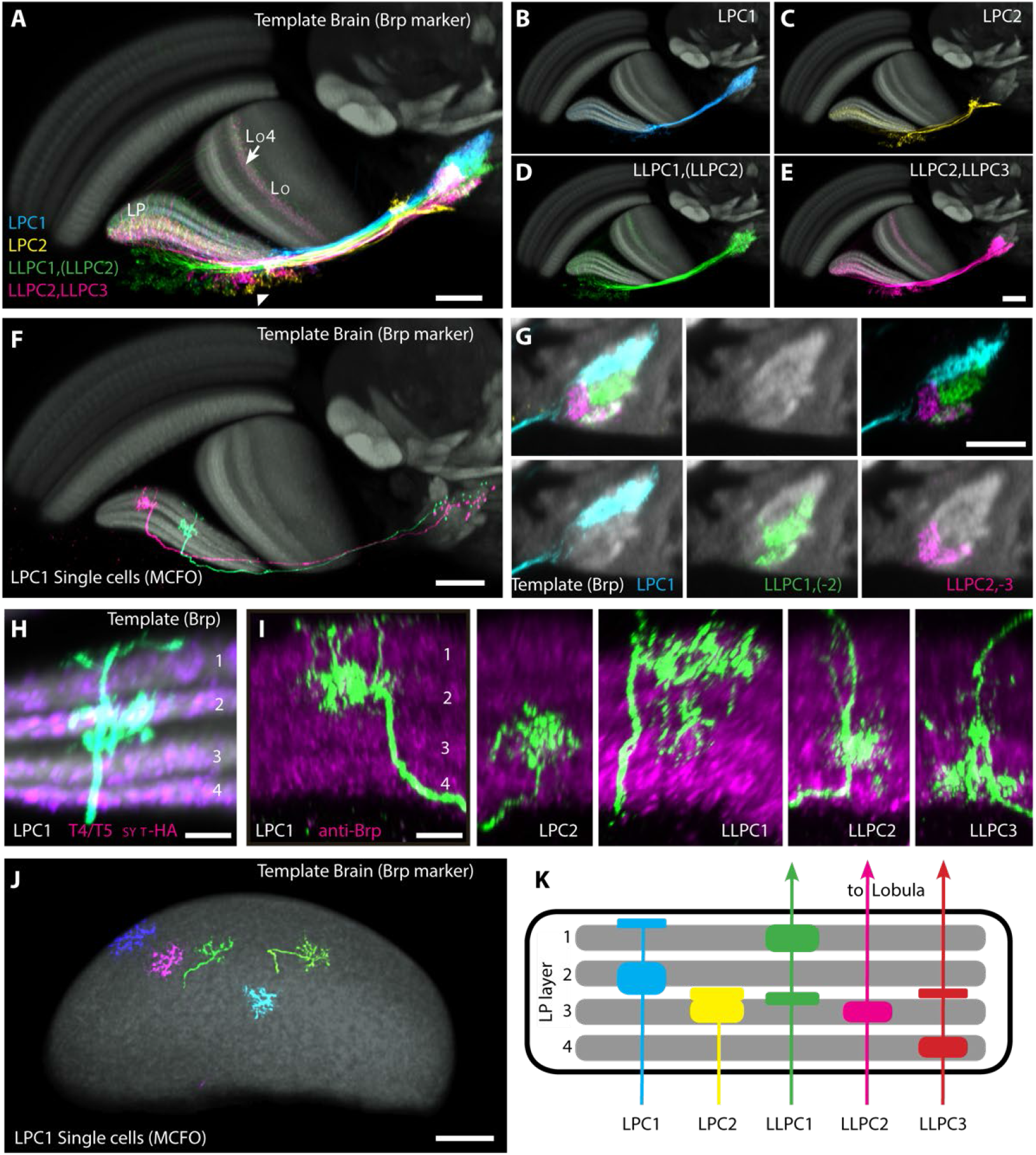
Small-field visual projections neurons of the lobula plate. (A-E) Projection patterns of <u>L</u>obula <u>P</u>late <u>C</u>olumnar (LPC) and <u>L</u>obula <u>L</u>obula <u>P</u>late <u>C</u>olumnar (LLPC) neurons. LPC (A,B,C) and LLPC (A,D,E) neurons have cell bodies in the lobula plate cell body rind, layer-specific arborizations in the lobula plate and project to the LPC glomerulus, a distinct, synapse-rich neuropile subregion in the posterior lateral protocerebrum (PLP). The LPC glomerulus is located posterior and ventral to most other optic glomeruli. LLPC cells also send a process to lobula layer Lo4. Images are composites of registered brains expressing a membrane marker in the indicated cell types and the neuropile marker of the template brain used for alignment. Anterior is up and medial to the left. Additional details of individual Figure panels, such as fly genotypes, can be found in Table 1. (F) Two individual LPC1 cells were labelled using MCFO^65^ and displayed as in (A-E). Single cells of the other cell types are shown in Fig. S2A-D. (G) Projections patterns of LPC and LLPC cell types reveal subdivisions of the LPC glomerulus. Single slices of composite image stacks of registered brains with a presynaptic marker (synaptotagmin-HA) expressed in the indicated cell types. Neuropile marker of the template brain in grey. (H) Sublayer structure of the lobula plate. Four synapse rich sublayers that contain presynaptic sites of directionally-selective T4/T5 neurons are separated by sublayers with lower synapse density. LPC1 processes in LP layer 2 overlap with the presynaptic sites of T4/T5 neurons in this layer. Images are composites of registered brains as above. The LPC1 cell (green) is one of the two neurons shown in (F). T4/T5 presynaptic sites were labeled by expression of synaptotagmin-HA in these cells. The Brp-based neuropile label of the template brain (in grey) provides an indication of overall synapse density. (I) Lobula plate layer patterns of LPC and LLPC cell types. LP layers were identified by anti-Brp labeling (magenta) and the relative position of the four main layers. (J) Spread of individual LPC1 cells in the lobula plate. Composite of MCFO labeled LPC1 cells from different registered image stack together with neuropile marker of the template brain are shown. Only signal within or very close to the lobula plate was included in the projections in the images. For some cells (here the two green neurons), this includes parts of neurites extending along the surface of the lobula plate. (K) Schematic illustrating the distinct optic lobe layer patterns of the five LPC and LLPC lobula plate VPNs distinguished in this study. For each cell type, the main overlap with T4/T5 terminals is in a single lobula plate layer (corroborated by EM connectome^18^). All cell types, especially LPC1, LLPC1 and LLPC3, also have branches in other lobula plate sublayers. LLPC cells also project to lobula layer Lo4. Scale bar in A, E, F, G, J is 20 µm and 5 µm in H, I.

Two of these cell types, called LPC1 and LPC2 (for Lobula Plate Columnar type 1 and type 2), innervate the lobula plate but not the lobula, while the other three, referred to as LLPC1, LLPC2 and LLPC3 (Lobula-Lobula Plate Types 1,2, and 3), send processes to the lobula in addition to the lobula plate (Fig. 2A-E, Fig. S2A-D). All five types share several anatomical features that distinguish them from other lobula and lobula plate projection neurons: all are small-field neurons, innervate specific lobula plate layers^18^, have cell bodies in the lobula plate cell body rind, and send axons from the posterior side of the lobula plate to the same distinct neuropile structure in the PLP which we refer to as the LPC glomerulus (Fig. 2A-E, Fig. S2A-D). Each cell type forms a population of ∼100 near-isomorphic neurons on each side of the brain. Dendrites of individual LPC and LLPC neurons span at most 10% of the retinotopic extent of the lobula plate (Fig. 2F,J, Fig. S2A-H). From this, we estimate these neurons to be sensitive to visual motion within a ∼20°-wide receptive field. Cell-type specific features of these neurons include the position of their terminals within the LPC glomerulus (Fig. 2A,G), the presence or absence of a lobula branch (Fig. 2A-E) and the innervation of distinct lobula plate layers (Fig. 2H,I,K). Since lobula plate layers are defined by the axon terminals of T4/T5 subtypes that each encode one of four directions of motion^16, 17, 19^ (Fig. 2H), the distinct layer patterns of each LPC and LLPC cell type (Fig. 2K) and recent connectivity data^18^, suggest that each subtype may selectively encode motion in a specific direction. For example, LPC1 cells primarily overlap and receive synaptic inputs^18^ from layer 2 T4/T5 neurons (Fig. 2H,I,K), which encode back-to-front motion.

To test the functional predictions of these anatomical findings, we developed split-GAL4^52, 53^ genetic driver lines for selectively targeting LPC and LLPC cell type (Fig. 3A, Fig. S3). These LPC and LLPC split-GAL4 lines were used to confirm and extend our anatomical analysis. To examine their behavioral roles, we crossed each split-GAL4 lines to *UAS-Shibire^ts^*^1^ and ran the F1 offspring through an extended protocol in the Fly Vision Box (see Methods). We found that LPC1-silenced flies showed significant, strongly diminished visual motion responses, while no other driver lines yielded substantial deficits across different stimulus conditions, although LPC2 silencing resulted in somewhat enhanced responses to visual motion (Fig. 3B). Using simultaneous expression of both Shibire^ts1^ and Kir2.1, which is expected to yield a more complete blockade of neuronal function by interfering with both synaptic transmission and membrane excitability, we obtained significant, strong deficits in the motion following behaviors across a range of stimulus conditions, for three different LPC1 split-GAL4 driver lines (Fig. S3A,B). Overall, these anatomical and behavioral results identify LPC1 neurons as disproportionately contributing to the Behavior-Anatomy Correlation Map’s hotspot in the PLP (Fig. 1E,G; hereon identified as the LPC glomerulus). Based on a recent transcriptomic study of many optic lobe cell types^54^, LPC1 neurons are expected to provide cholinergic input to their targets in the glomerulus, suggesting they form an important, excitatory pathway for relaying visual motion information to the central brain.

**Fig. 3.**
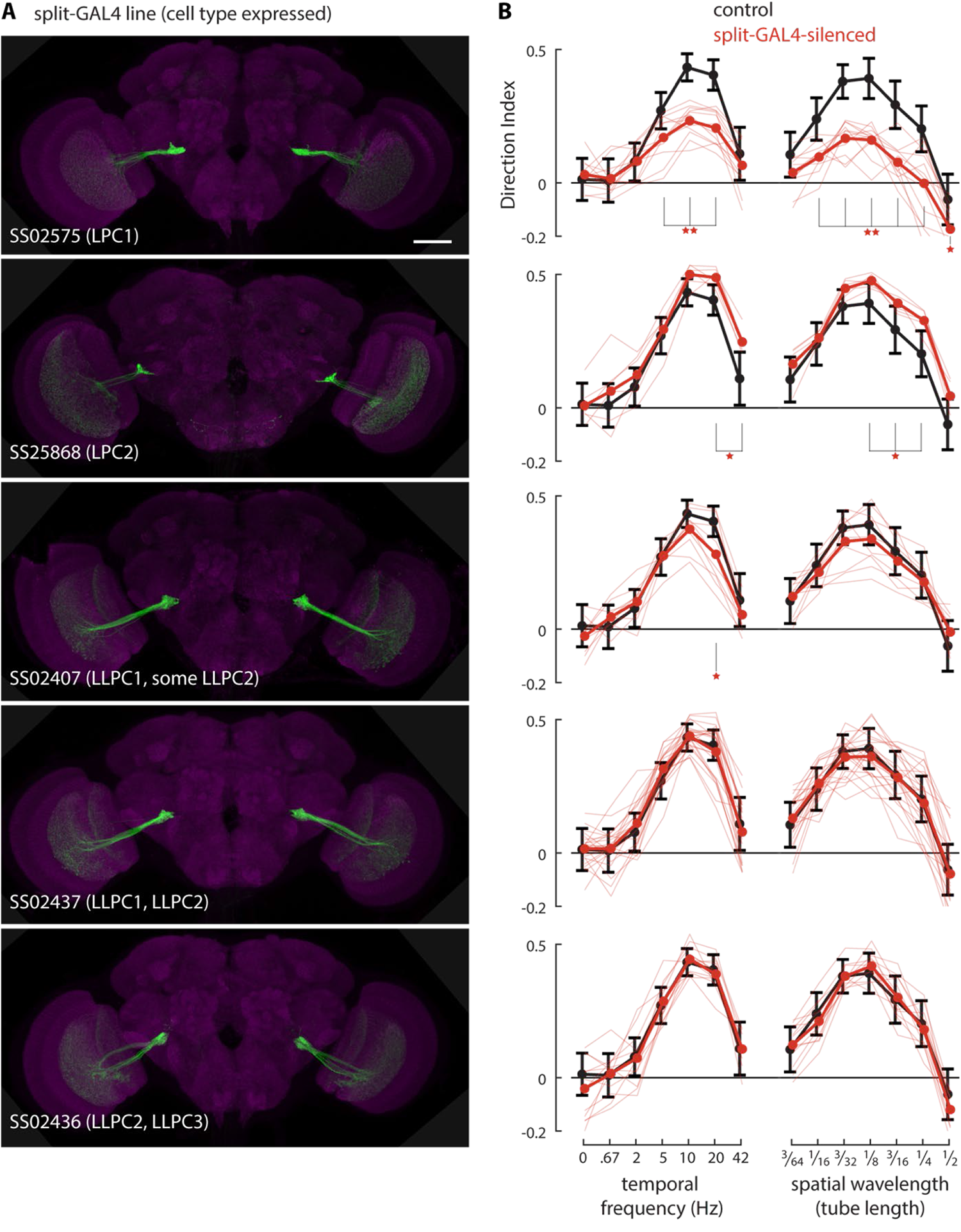
LPC1 neurons contribute to following visual motion. Split-GAL4 lines were developed to target expression to the small-field Lobula Plate Projection neurons. The expression pattern images (A) are maximum intensity projections through the full depth of fly brains. Each indicated split-GAL4 driver was used to express a membrane marker, shown in green, while a neuropile marker (anti-Brp) is shown in magenta. Scale bar, 50 µm. The effect of silencing the neurons (by expressing shibire^ts1^) on visual motion following in the Fly Vision Box is shown in (B), alongside the corresponding expression patterns in A. Visual motion responses from an extended protocol in the Fly Vision Box to gratings moving at a range of temporal frequencies and spatial wavelength (of the grating cycle, all moving with a temporal frequency of ∼10 Hz), summarized as tuning curves, quantified with a Direction Index (see Methods). The behavior of each split-GAL4 line > UAS-Shibire^ts1^ is plotted in red, with individual tube data shows in lighter lines (6-24 tubes per genotype), and the mean across tubes shown in thicker lines, and is compared to the split-GAL4 control genotype (in black, mean ± s.d. of 208 tubes of flies from 36 experiments. Silencing LPC1 neurons leads to large, significant reductions in following visual motion, a result further confirmed by ‘doubly silencing’ these cells using several LPC1-expressing split-GAL4 lines (Fig. S1). Statistical significance between experimental and control genotypes determined using Mann-Whitney U-test controlled for False Discovery Rate: 1 star = p < 0.05; 2 stars = p < 0.01. Additional images of the split-GAL4 lines are in Fig. S3B,C. The genotypes contributing to this Figure are in Table 1.

### LPC1 neurons play a critical role in the behavioral response to back-to-front translational motion

Flies walking in the Vision Box will experience some rotational optic flow, especially when they are changing direction (turning), but their dominant visual experience will be of translational optic flow (Fig. 4A). Because of this, the effect of silencing LPC1 neurons could either be due to a deficiency in the visual guidance of re-orientation or of forward walking, or both. In order to uncover how LPC1 neurons contribute to reduced following of visual motion (Fig. 3A, Fig. S3A), we examined the behavior of individual tethered flies walking on an air supported ball^55^ responding to visual stimuli presented on a cylindrical LED display^56^. In contrast to the Fly Vision Box, the tethered fly experiments allow precise control over the visual experience of each fly together with higher resolution measurements of walking behaviors.

**Fig. 4.**
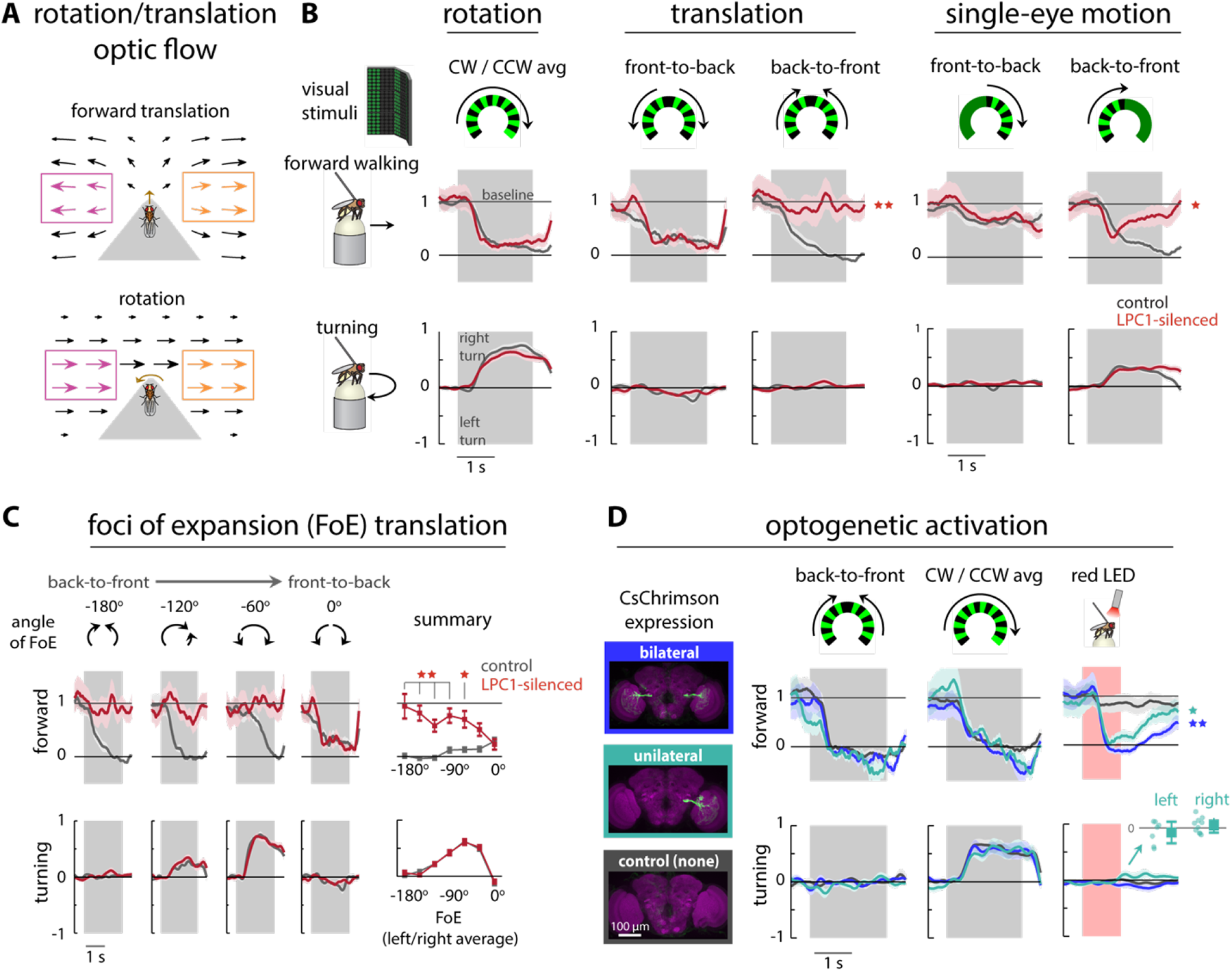
LPC1 neurons play a critical role in the behavioral response to back-to-front translational motion. (A) Illustration of optic flow experienced by a fly moving forward (translation) or turning to the left (rotation). Arrows plotted in Mercator projection (cosine corrected) with size of arrows proportional to the apparent angular speed. (B) Forward walking and turning responses of tethered flies walking on an air-supported ball to moving visual grating patterns. Flies were positioned in the center of the cylindrical LED display. The visual stimuli present either rotational or translational visual motion, or motion restricted to one eye, and are schematized using a top-down view of the arena. Mean time courses of the forward and turning components of the walking responses are plotted as mean ± SEM (shaded) for n = 20 LPC1-silenced flies in red and n=20 genetic control flies in black. Grating patterns (30° spatial wavelength) moved with 9 Hz temporal frequency corresponding to 270°/s (responses to additional speeds in Fig. S4). Grey box indicates when moving gratings are presented, otherwise the flies are presented a uniform (blank) pattern of intermediate brightness. (C) Left: Mean time course of turning and forward components of the tethered walking responses to translational visual motion presented with different foci of expansion. The data are plotted as in B. Right: The summarized forward walking and turning responses to the complete set of experimental conditions with translational motion emanating from foci of expansion located every 30° around the fly. (D) Left: Examples of fly brains where CsChrimson was stochastically targeted to LPC1 neurons (using established method^47^). Flies were dissected and imaged after behavioral experiments, so flies could be grouped based on expression: n=19 flies with bilateral LPC1 expression, dark blue; n=17 flies with unilateral expression, in teal; n=14 flies lacking expression that serve as the negative control, in black; example maximum-intensity projections of confocal stacks shown on the left. Right: Mean time course of forward and turning components of walking responses to back-to-front translational motion, rotational motion, and optogenetic activation with a fiber-coupled red LED. Bottom-right inset: Mean turning response of flies with unilateral CsChrimson expression separated into left or right hemisphere expression (n=7 left, n=10 right). Statistical significance between experimental and control flies determined using Mann-Whitney U-test controlled for False Discovery Rate: 1 star = p < 0.05; 2 stars = p < 0.01. Turning responses are presented for one direction but are averaged across L/R symmetric conditions (see Methods). The genotypes contributing to this Figure are in Table 1.

When large-field rotational visual motion was presented to tethered walking flies (‘empty-split’ control genotype), they slowed down while simultaneously turning in the direction of the rotating motion (Fig. 4B, left; summary for multiple speeds and results of statistical tests in Fig. S4), exhibiting the stabilizing reaction referred to as the optomotor response^4^. In comparison to control flies, silencing LPC1 neurons had no effect on either the turning or the slowing down components of the rotational optomotor response. Considering the strong effect of silencing LPC1 on visual motion following in the transparent corridors of the Fly Vision Box, we reasoned that perhaps a different ‘optomotor’ reaction was relevant for this visual condition, one that is downstream of the detection of visual translation as opposed to visual rotation. To investigate this possibility, we exposed flies to the simplest visual stimuli that approximate forward or backward translational optic flow—two opposing halves of a rotational optic flow field (either front-to-back or back-to-front) for each eye. These right-left symmetric stimuli did not result in consistent directional turning, but both caused control flies to reduce their forward walking (in agreement with recent work^33^). Remarkably, the reduction in forward walking in response to fast back-to-front, but not front-to-back, motion was abolished when the LPC1 neurons were silenced (Fig. 4B and Fig. S4).

To better understand how the organization of optic flow contributes to the effect of silencing LPC1 neurons, we divided these stimuli into their constituent left and right halves and presented them individually to tethered flies. In response to monocular back-to-front and front-to-back motion (Fig. 4B, right), control flies slowed down and turned (in the direction of motion, larger for the back-to-front stimulus; other speeds in Fig. S4). The flies with LPC1 silenced exhibited a significantly reduced slowdown in response to back-to-front motion, while their other behavioral reactions, including turning to this same stimulus, and walking and turning to front-to-back motion, were unaffected (Fig. S4, right). This result suggests that LPC1 neurons mediate behavioral reactions specifically to back-to-front visual motion, and that the behavioral consequence of LPC1 activity may primarily contribute to the regulation of forward walking speed, and not to the body turning that is also evoked by the same visual stimulus.

To further probe the specificity of the LPC1-silencing effect, we examined responses to a set of translational optic flow fields that correspond to cases when the direction of gaze is not aligned with the direction of translation (looking in one direction while moving in another, recently examined in the context of coordinate transformations in the central complex^57^). To simulate this visual condition, we presented a panoramic pattern consisting of motion emanating from a Focus of Expansion (FoE), positioned at multiple azimuthal locations. In response to this panoramic translational motion (several examples in Fig. 4C), control flies slowed down and turned away from the FoE (turning behavior remarkably similar to that measured in flying flies^31^). The effects of silencing LPC1 are strikingly specific, with a significant reduction in the slowing down reaction, but without any effect on turning. Moreover, the lack of slowing down in LPC1-silenced flies is most prominent in the conditions where large parts of the display showed back-to-front motion (Fig. 4C, summary). Across this comparative set of visual stimuli, these results clearly demonstrate that silencing LPC1 neurons leads to a specific deficit in forward walking, but not turning, in response to back-to-front translational, but not rotational, visual motion.

While these silencing effects suggest a simple causal link between the perception of back-to-front visual motion and the control of forward walking, they do not rule out other scenarios. One such possibility is that LPC1 cells, in addition to other neurons, also contribute to turning behaviors, and there is sufficient redundancy and/or complexity in these pathways to mask the effects of LPC1 silencing. To address this possibility, we set out to isolate the contribution of LPC1 neurons to walking behaviors using optogenetic activation. However, the synchronous and simultaneous depolarizations of LPC1 neurons from both eyes could again mask a contribution to turning behaviors. To avoid this confound, we used a previously developed stochastic genetic method^47^ to target CsChrimson^58^ to LPC1 neurons on only one side of the brain. To determine the expression pattern of each individual fly, we dissected and imaged each animal after the behavioral experiment. Because this method produces animals with bilateral as well as no expression, the individuals with these genotypes served as positive and negative controls. We first wanted to confirm that these flies had similar behavioral reactions to rotational and translational optic flow, and indeed all three groups of flies (those with either bilateral, unilateral, or no LPC1 CsChrimson expression) responded similarly (Fig. 4D), and similarly to our control flies for the silencing experiment (Fig. 4B). To activate CsChrimson with minimal effects on vision, we used a 660 nm LED coupled to a fiber directed at the head of each tethered fly. In response to this optogenetic depolarization of LPC1 neurons on both sides, flies reduced their forward walking (to an approximate stop) without any effect on turning behavior (Fig. 4D, right). Flies lacking CsChrimson expression did not slow down or turn in response to optogenetic activation, whereas flies with unilateral CsChrimson expression slowed down nearly as much as those with bilateral CsChrimson expression. While the population average of the unilateral CsChrimson flies did not exhibit an LPC1-induced turning behavior, we were careful to note which side of the brain contained LPC1 neurons expressing CsChrimson. Even when separated by the side of expression (Fig. 4D, right, inset), we find that unilateral LPC1 activation did not evoke a directional turning bias. These activation results clarify the contribution of LPC1 neurons to behavioral control. The bilateral activation of these excitatory neurons, as would be experienced by moving backwards relative to a static environment, leads to a slowing down of forward walking. Activation of these neurons, even on one side of the animal, also leads to slowing down, but not to turning, consistent with a specific link between back-to-front motion detection and the control of walking speed.

### LPC1 neurons selectivity encode back-to-front translational visual motion by integrating contralateral motion

Visual projections neurons that bridge the optic lobe and the central brain provide a unique opportunity to understand how selectivity for specific visual features maps onto behavioral control. Do LPC1 neurons encode back-to-front visual motion as is suggested by their anatomy (Fig. 2I), connectivity^18^, and our behavioral analysis (Fig. 4B,C)? To determine the visual motion selectivity of LPC1 neurons, we performed *in vivo* two-photon calcium imaging of the LPC1 population axon terminal bundle (in the glomerulus) while presenting visual motion stimuli (Fig. 5A). In this experimental setup, the fly’s head is pitched down, allowing access to the posterior side of the central brain of flies with GCaMP6m^59^ expressed in LPC1 neurons. A recently designed, higher resolution, cylindrical LED arena^41^ was positioned such that it was aligned to the equator^60^ of the fly’s eye (Fig. S5A,B). All calcium activity measurements were from the left LPC1 population, expected to convey ipsilateral visual motion from the left eye.

**Fig. 5.**
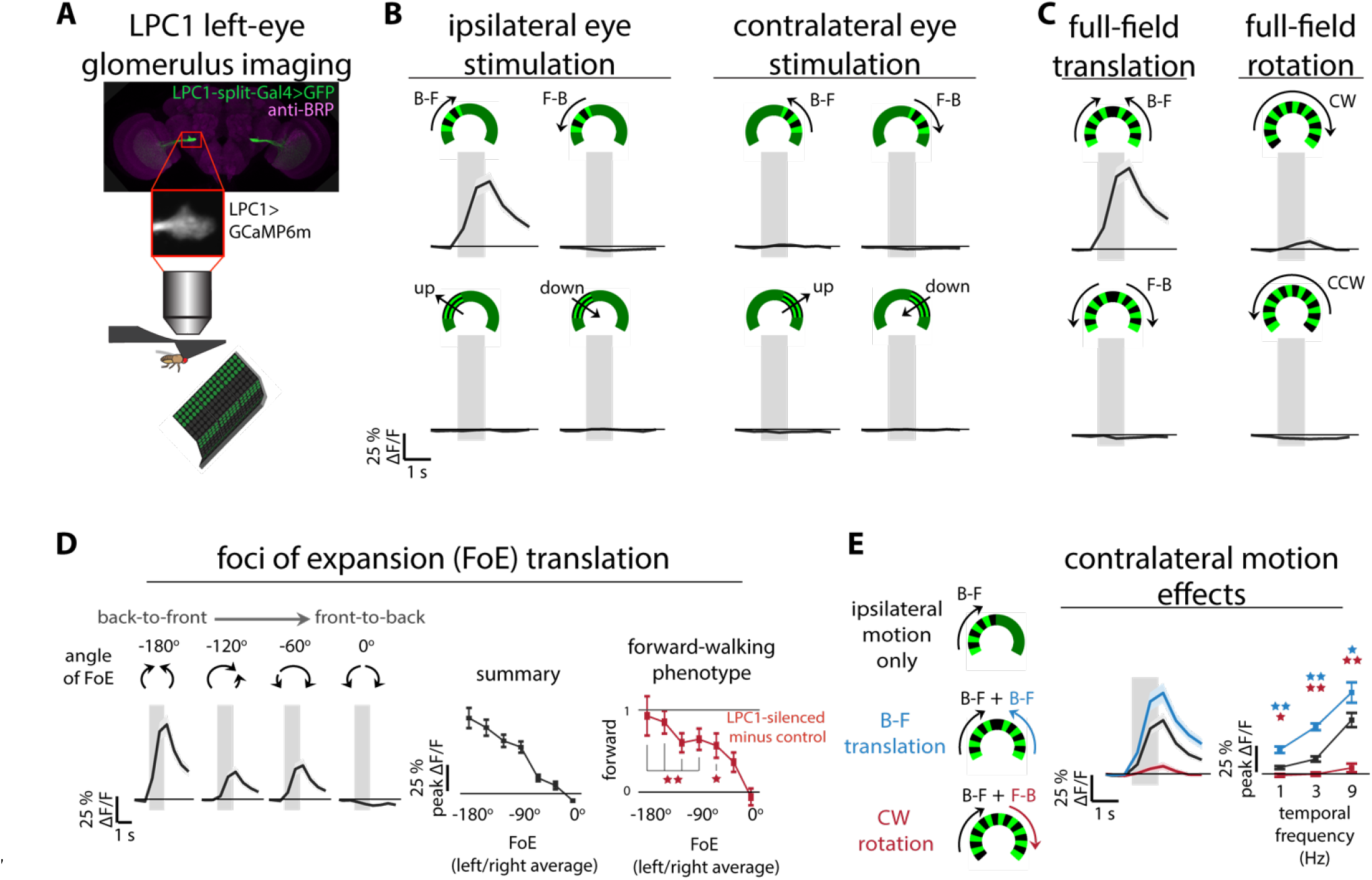
LPC1 neurons selectively encode back-to-front translational visual motion by integrating contralateral motion. (A) Schematic representation of *in vivo* two-photon calcium imaging setup. The activity of the population of LPC1 neurons on the left side, expressing GCaMP6m, is imaged from the glomerulus, while the fly views visual stimuli. (B-E) The time course of the left LPC1 glomerulus calcium response to visual grating motion (30° spatial wavelength, moving at 9 Hz temporal frequency, presentation interval indicated with the gray bar). Stimuli are schematized showing the top-down view of the display. Data displayed as mean ± SEM (shaded region) for n=10 flies. (B) LPC1 responses to grating stimuli moving in four directions, presented to the left and right eyes independently. The stimulation window spanned 20-90° in azimuth measured from the midline of the fly’s visual field. (C) LPC1 responses to full-field rotational and translational motion spanning -90° to 90° in azimuth. (D) Left: LPC1 responses to translational motion presented with the Focus of Expansion (FoE) at different azimuthal positions. Right: Summary of the peak (± SEM) LPC1 calcium responses to the translational stimuli emanating from different foci of expansion, plotted alongside the LPC1 silencing behavioral phenotype (data from Fig. 4C re-plotted as the difference in forward-walking between LPC1-silenced flies and control flies). (E) Mean (± SEM) time course of LPC1 calcium responses to ipsilateral back-to-front motion presented alone or paired with contralateral back-to-front and front-to-back motion. The responses across several stimulus speeds are shown summarized on the right and responses to related stimuli shown in Fig. S5C,D. Statistical significance between the bilateral and the ipsilateral responses is assessed using the Mann-Whitney U-test corrected for False Discovery Rate: 1 star = p<0.05; 2 stars = p<0.01. All calcium imaging data in this figure come from the same set of n=10 flies.

We compared the response to moving gratings presented to the left and right eye separately, in all four cardinal directions. The only detectable responses from the LPC1 glomerulus population were to back-to-front visual motion presented to the left (ipsilateral) eye (Fig. 5B), consistent with the anatomical and behavioral predictions. However, the behavioral data does not make a strong prediction about whether LPC1 neurons might integrate bilateral motion information. The large effect of silencing LPC1 neurons on bilateral back-to-front motion responses (Fig. 4B,C) could be attributed to downstream neurons that integrate from both LPC1 populations. To see whether bilateral information is combined in LPC1 neurons, we presented rotational and translation motion stimuli spanning both eyes. As expected, the left LPC1 glomerulus responded strongly to back-to-front bilateral visual motion, but not to front-to-back visual motion (Fig. 5C). Surprisingly, the response to rotational visual motion containing back-to-front visual motion on the left eye showed a significantly reduced response, suggesting that the global optic flow pattern influences the response of LPC1 neurons (Fig. 5C). Furthermore, the calcium responses of LPC1 neurons to translational patterns presented with a focus of expansion (FoE) at multiple positions around the azimuth of the fly’s visual field confirm their maximal selectivity for back-to-front visual motion, with reduced activation for other positions of the FoE (Fig. 5D, left). Remarkably, this graded response is very similar to the effect of silencing LPC1 neurons on forward walking, in response to the same stimuli (Fig. 5D, right, comparison to behavioral data adapted from Fig. 4C). Thus, we find LPC1 neurons encode the precise visual stimuli for which we have also shown them to be behaviorally required.

The large difference between the LPC1 calcium responses to presentations of translational and rotational optic flow that both contain the same amount of ipsilateral back-to-front motion (Fig. 5C), suggests that contralateral visual motion modulates the response to ipsilateral motion – providing global ‘context’ to an otherwise ambiguous visual scene. To probe this mechanism further, we presented back-to-front ipsilateral visual motion (Fig. 5E, in black) paired with either contralateral back-to-front motion (mimicking translation, in blue) or front-to-back motion (mimicking rotation, in red). We note that these contralateral motion stimuli, presented on their own, did not evoke a detectable calcium response (Fig. 5B), and yet these contralateral visual inputs modified the response to ipsilateral back-to-front motion (Fig. 5E). LPC1 responses were significantly enhanced for translation-like stimuli and were significantly reduced for rotational stimuli, at all tested speeds (Fig. 5E). We further confirmed that this result holds whether the region of frontal binocular overlap is stimulated or not (Fig. S5C), and that LPC1 neurons encode translational optic flow regardless of its fine structure (Fig. S5D). Taken together, these experiments demonstrate that the LPC1 population detects back-to-front visual motion and integrates contralateral visual motion information to enhance their selectively for translational, as opposed to rotational, optic flow.

### An additional small-field neuron type, LLPC1, encodes translational visual motion with directional selectivity opposite to LPC1 neurons

After discovering LPC1’s strong preference for translational, back-to-front optic flow, we wondered whether other small field lobula plate neurons might show similar preference for optic flow patterns that accompany other types of translational motion. We then examined the response properties of the Lobula-Lobula Plate Columnar Type 1 cells (LLPC1), which receive their primary inputs in layer 1 of the lobula plate^18^ (Fig. 2I,K) and are therefore expected to encode the opposite direction of visual motion as LPC1. As with LPC1, their anatomy does not suggest a specific role in the detection of either translational or rotational visual motion. Perhaps surprisingly, silencing the LLPC1 neurons did not yield any statistically significant behavioral phenotype in either the Fly Vision Box (Fig. 3A) or in the fly-on-ball walking assay (Fig. S6).

We imaged the calcium responses of LLPC1 neurons in the glomerulus, following the methods and stimulus protocol developed for LPC1 neurons (Fig. 6A). As predicted by their anatomy, and despite their lack of a strong silencing phenotype, the LLPC1 neurons were selectively activated by ipsilateral front-to-back motion but not by contralateral visual motion (Fig. 6B). We noted a modest response to upward motion (Fig. 6B); this response is most likely a contribution of LLPC2 expression within our LLPC1-split-GAL4 line. LLPC2 axons terminate adjacent to the LLPC1 axons (Fig. 2G), and further analysis (Fig. S7A-B) showed that the upward motion response is restricted to a glomerulus region that is distinct from the front-to-back responses. We thereby have also inadvertently established that LLPC2 neurons, with inputs in layer 3 of the lobula plate^18^ (Fig. 2I,K), have the expected property of encoding upwards visual motion.

**Fig. 6.**
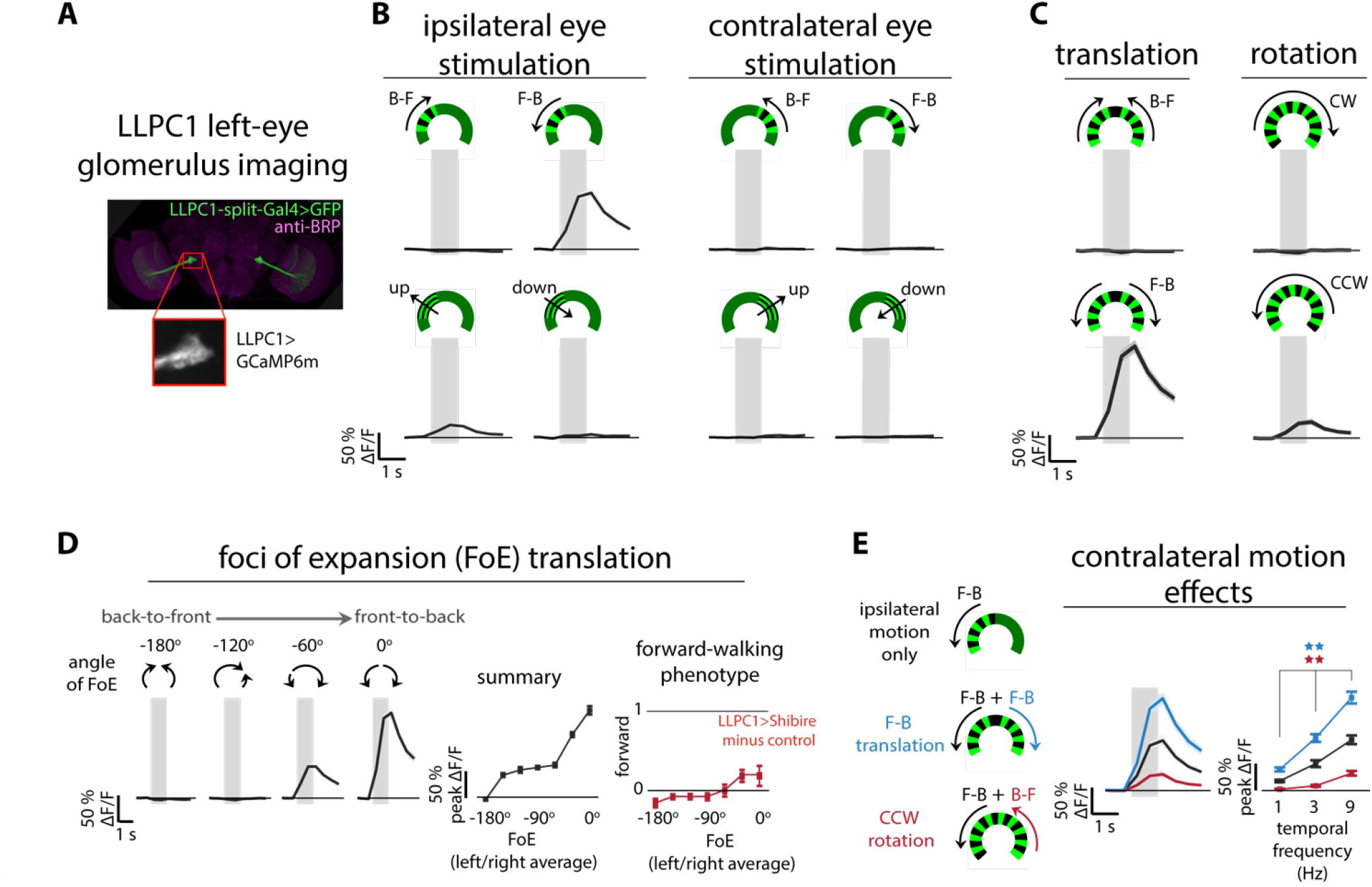
An additional small-field neuron type, LLPC1, encodes translational visual motion with directional selectivity opposite to LPC1 neurons. (A) The visual motion evoked calcium responses of the left side LLPC1 neurons, expressing GCaMP6m, were obtained by imaging from the glomerulus, following the methods of Fig. 5. (B-E) Time course of the left LLPC1 population calcium response to the identical visual motion stimulus protocol used for the functional imaging of LPC1 neurons in Fig. 5. The calcium imaging data are based on n=10 flies and are plotted using the same format and conventions as Fig. 5, where the most noteworthy difference is the direction of preferred motion—LLPC1 neurons responds primarily to front-to-back visual motion (but see Fig. S7A,B for further details on the modest response to upwards motion). In (D) the responses to translational motion emanating from different foci of expansion are compared to the behavioral results of silencing LLPC1 neurons (behavioral data from Fig. S6B re-plotted as the difference in forward-walking between LLPC1-silenced flies and control flies). (E) Contralateral front-to-back motion enhances the ipsilateral response, while contralateral back-to-front motion suppresses it, consistent with selective tuning for forward translational motion (responses excluding binocular overlap zone in Fig. S7C). Statistical significance is assessed using the Mann-Whitney U-test corrected for False Discovery Rate: 1 star = p<0.05; 2 stars = p<0.01.

The responses of the LLPC1 neurons to optic flow patterns containing translational visual motion are remarkably consistent with the LPC1 responses, except that the neurons prefer front-to-back translational visual motion (Fig. 6C). As with LPC1, the neurons showed a graded response to translational visual motion presented at different FoE positions, with the peak response occurring for front-to-back motion (Fig. 6D). Comparing the visual responses of the LLPC1 neuron to the behavioral effect of silencing this cell type, we find a similar trend, but the LLPC1 phenotypes do not rise to the level of statistical significance. We therefore cannot rule out either that the silencing effect is incomplete in these cells, or the parallel contribution of additional cell types contributing front-to-back visual motion to the control of forward walking.

Finally, we found a similar contribution of contralateral visual motion to establishing the sensitivity for translational visual motion. LLPC1 neurons’ response to ipsilateral front-to-back visual motion was enhanced by contralateral front-to-back motion and suppressed by contralateral back-to-front motion for all tested speeds (Figs. 6E, S7C). Front-to-back visual motion is a ubiquitous feature of navigation through any environment and these results establish LLPC1 neurons as an important sensory pathway for detecting the visual consequence of forward self-motion together with a mechanism for discarding confounding visual motion during turning.

### Cascaded inhibition from neurons tuned to opposite directional motion establishes LPC1’s selectivity for translation motion

The ‘global context’ provided by contralateral visual motion modulates the responses of LPC1 and LLPC1 to their ipsilateral visual input, promoting the selectivity for translational optic flow (Fig. 5E, 6E). A mechanistic explanation for this selectivity would be that these cell types receive inhibition from neurons encoding contralateral non-preferred motion, which would explain why we did not see any responses to contralateral motion alone (calcium indicators are often unreliable reporters of hyperpolarization). Furthermore, the enhancement we observed when contralateral preferred direction motion was also presented (back-to-front for LPC1 and front-to-back for LLPC1) could be the excitatory contribution of additional cell types (too weak to be observed in isolation) or a consequence of reducing the contralateral inhibition.

This inhibition due to contralateral motion could arise from synaptic inputs onto LPC1 and LLPC1 cells either in their dendrites in the lobula plate or their axon terminals in the glomeruli (by separate midline-crossing neurons for each cell type). The lobula plate houses layer-specific inhibitory neurons^18, 61^, but as these neurons would inhibit both the rotation and translation sensitive outputs of the lobula plate, these seem unlikely to serve a role in promoting translational optic flow sensitivity. We instead focused on the axon terminals, which could provide a compact target for the coordinated inhibition of the entire population of small field neurons. Before the availability of the hemibrain dataset, we undertook a neuroanatomical exploration of neurons that appear to provide input to the LPC1 glomerulus and examined many candidate cell types. The most promising cell type we identified in this search is a large neuron, present once per central brain hemisphere, with extensive arbors in both the LPC1 and LLPC1 subregions of the LPC glomerulus (named LPC-IN; Fig. 7A). Synaptic marker expression suggests that this cell is presynaptic in the LPC1 glomerulus and postsynaptic in the LLPC1 glomerulus and based on the expression of GAD1, is GABAergic, and thus likely inhibitory (Fig. 7B,C,D). This single neuron can be unambiguously matched to a cell named PLP249 in the hemibrain^36^, and the connectivity data for this cell confirms our predictions for substantial inputs from LLPC1 neurons and substantial outputs onto LPC1 neurons. LPC-IN is expected to inhibit LPC1s. To test this, we developed a split-GAL4 line to target LPC-IN and imaged calcium responses to visual stimuli. We found that LPC-IN primarily encodes ipsilateral front-to-back motion (Fig. 7B). However, it does not respond to contralateral visual motion and is thus unlikely to be a source of inhibition due to contralateral visual motion. The anatomy and response properties of this input neuron suggest it inherits its (front-to-back) visual responses from LLPC1. Lacking the optic lobes and the contralateral brain hemisphere, the hemibrain data set is a challenging resource in which to identify this contralateral inhibition pathway, but future EM resources should reveal this complete circuit. Nevertheless, the identification of this new cell type establishes another example of the use of inhibition by oppositely tuned neurons for the generation of response selectivity along the motion pathway—but now well outside of the visual system (Fig. 7F).

**Fig. 7.**
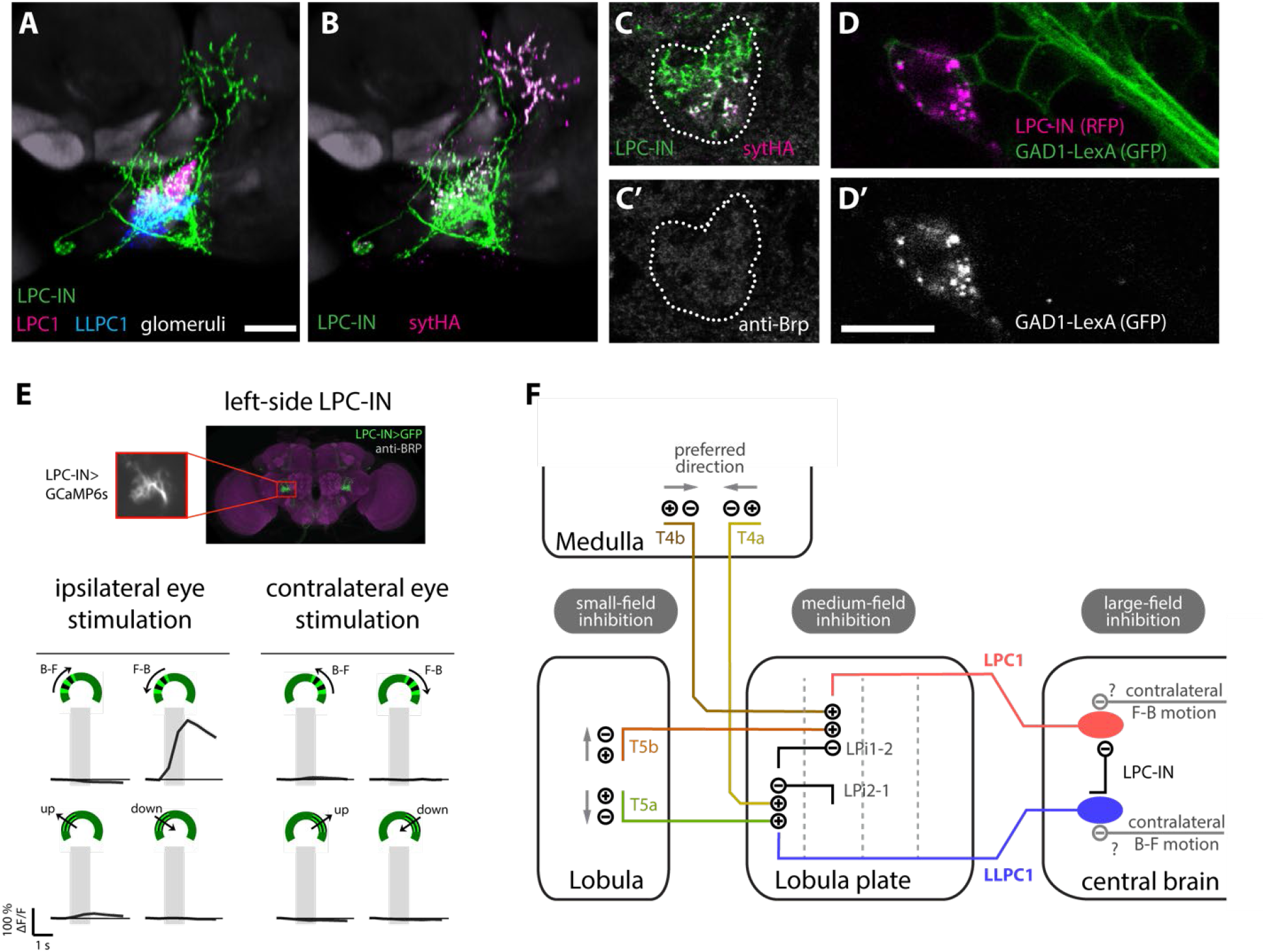
Cascaded inhibition from neurons tuned to opposite directional motion establishes LPC1’s selectivity for translational motion. (A) Overlay of registered brains (same orientation as Fig. 2A-G) showing an interneuron, LPC-IN (labeled with a membrane marker; green). The LPC1 and LLPC1 glomeruli are indicated in cyan and magenta, respectively. Processes of LPC-IN are found throughout both the LPC1 and LLPC1 target regions. The neuropile label of the reference brain is in grey. Scale bar, 10 µm. (B) Overlay as in (A) with a pre-synaptic marker (syt-HA) shown in magenta. (C,C’) Distribution of the syt-HA marker in a single section through the LPC glomerulus. Based on comparison with registered samples (as in A,B) most LPC-IN presynaptic sites within the LPC glomerulus overlap with LPC1, not LLPC1. (D,D’) LPC-IN expresses GAD1. Double labeling showing LPC-IN (magenta) and a GAD1-LexA reporter (green in D, grey in D’). FISH against GAD1, ChAT and VGlut (see Fig. S8) also supports a GABAergic phenotype of LPC-IN. Scale bar, 10 µm. (E) *In vivo* two-photon calcium imaging of the left side LPC-IN, using a split-GAL4 line expressing GCaMP6s, in response to visual motion stimuli. The mean (± SEM) time course of the left LPC-IN calcium responses to the identical stimulus protocol as in Fig. 5B, 6B, from n=4 flies. (F) Summary diagram of the direction-selective circuitry contributing to LPC1 and LLPC1 selectivity for translational motion in both the optic lobe and central brain, emphasizing the importance of inhibition. Directional preferences are first established by the T4 and T5 neurons, where small-field, trailing-side inhibition plays a critical role^10, 15, 66^. In the lobula plate, the bi-layer LPi neurons sharpen the directional tuning inherited from T4/T5 neurons via motion opponency^61^ acting on LPC1 and LLPC1^18^. In the central brain, the LPC-IN cell reinforces the directional sensitivity established in the optic lobe by inhibiting LPC1 neurons in responses to LLPC1 activation. The pathways for LPC1 and LLPC1 modulation by contralateral motion have not been identified but are hypothesized to be acting on the glomeruli.

## DISCUSSION

Motion processing in flies is a well-studied field, and yet the primary pathways connecting the processing of optic flow^1, 3^, large patterns of visual motion induced by movement, to the control of locomotion have not been delineated. Most previous studies have focused on the wide-field Lobula Plate Tangential Neurons, many of which respond best to rotational visual motion^24, 28, 62^. In this study we used a systematic analysis of visual motion response deficits in walking flies with subsets of neurons silenced, to identify a neuroanatomical locus with a critical contribution to visual motion behaviors (Fig. 1), and then described several cell types (Fig. 2) within this region. One of these small-field Lobula Plate Columnar cell types, LPC1, is required for regulating forward locomotion in response to backwards, translational visual motion (Fig. 3, 4). Further examination of LPC1 cells, along with another cell type, LLPC1 revealed that these neurons are specifically tuned to the optic flow a fly would experience while translating backwards and forwards, and that this selectivity for translation over rotation is established via contralateral inhibition (Fig. 5, 6, 7). Taken together, these results establish the small-field output neurons of the lobula plate as a major conduit for translational optic flow. It is likely that they feed into circuits controlling locomotion, which can now be systematically explored using recent connectome data including the central brain^36, 63^.

### Regulating forward locomotion using translational optic flow

We established the Fly Vision Box as a high-throughput assay suitable for measuring locomotion of flies in response to visual stimuli. In retrospect we realized that the primary experience of flies in this assay is of translational optic flow. This feature has proven to be fortuitous, since the pathways and neurons that encode translational visual motion—likely the dominant visual experience of seeing animals moving through the world—is not well known. While previous work in blowflies had identified LPTs with translation-like properties^34^ and mixed sensitivity for both translational and rotations motion in HS and VS cells^62^, our work presents the clearest demonstration of visual neurons specialized for detecting translational visual motion. Moreover, we found two cell types, LPC1 and LLPC1 that are selectively turned for translational body motion in either the forward or backward direction, and thus likely critical for the visual regulation of forward locomotion. Based on the population-level activity measurements we made of LPC1 neurons, they are not sensitive to the fine structure of translational motion, as evidenced by the similar responses to more geometrically accurate representations of this motion (Fig. S5D).

Finding a translation-sensitive system built from populations of LPC1s and LLPC1s, small ipsilateral neurons, is an unexpected result. Due to their small size, these cell types had been underexamined in electrophysiological studies of larger flies, and by virtue of their small field of view, they cannot individually encode a translational flow-field. Pioneering work on optic flow^3^ showed that accurate estimates of self-motion require large coverage of the visual field. This theoretical insight, as well as our results (Figs. 5,6,7), clarifies that the selectivity for translation over rotation is established by integrating motion signals across the two eyes. But what benefit do small neurons have for detecting translational body movements, a pattern with a global consequence? One suggestion from our data (Fig. 5D, 6D) is that these cells might be used to estimate the travelling direction of the animal in conditions where the direction of gaze is not aligned to the heading direction, resulting in non-frontal foci of expansion/contraction^31, 57, 64^. Continuing down this speculative path, the suggestion is that smaller neurons provide flexibility for encoding different patterns of optic flow, and perhaps for encoding the potentially large dynamic range of local motion speeds found within translational flow fields. The geometry of optic flow is such that nearby objects will induce much larger local motion during translation (but not rotational) motion^3^, and so it is possible that a population of small neurons is better able to represent the structure of translational optic flow fields, since individual neurons can more faithfully encode the range of motion strengths without saturating. A future simulation study should carefully explore the trade-offs implied by this speculative proposal.

It is noteworthy that while LPC1 silencing led to profound effects on forward walking, LLPC1 silencing did not produce a similar forward walking deficit in response to translational motion. This finding may suggest that the control of visual motion induced by forward walking, expected to be a critical component of many behavioral programs, is likely more complex. It is possible that LLPC1 neurons, whose responses are well-matched to the idealized visual stimuli accompanying forward locomotion, may not directly contribute to the control of walking. Similarly, the repertoire of neurons encoding this visual motion and contributing to the behavioral control might be larger than in the LPC1 circuit, and thus more resilient to silencing.

The simpler, causal role of LPC1 in regulating walking speed uncovered an unexpected result about locomotion control. We found that forward and turning components of walking behavior can be largely decoupled. When flies execute a rapid turn, they typically reduce their forward locomotion^5^ (and Figure 4B), but as we show, silencing LPC1 and presenting visual stimuli, especially the FoE stimulus in Fig. 4C, exposed a clear deficit in the slowing down in forward walking without effecting turning. This stimulus further revealed a stunning concordance between behavioral effects of silencing and the neurons’ visual sensitivity (Fig. 5D), suggesting that we have identified the critical transformation for this sensorimotor behavior—linking translational visual motion to the control of forward walking.

### Cascaded Inhibition as a prominent feature of the motion pathway

While the mechanism by which contralateral visual motion signals suppress rotational motion responses in LPC1 (and LLPC1) neurons remains to be established, the discovery of the LPC-IN underscores the critical role of inhibition by neurons tuned to opposing directions of motion in sharpening directional selectivity. This computation, termed motion opponency^51^, now appears to be a general feature of motion detection at all scales for which motion is processed in the fly brain (Fig. 7C). The integration of offset excitatory and inhibitory contributions to the T4 and T5 neurons is perhaps the first instance of opponency that is required for computing directional selectivity. These offset small-field inhibitory inputs correspond to 1-2 ommatidia, or <1% of the field of view of each eye^10^. In the lobula plate, the neurons integrating from the directionally selective neurons, such as the VS cells^61^ or the LPLC2 looming sensitive neurons^51^, receive inhibitory contributions from interneurons of the Lobula Plate (LPi cells), which themselves receive inputs from T4/T5 and produce a sign inverted opponent motion signals^18^. Different bi-layer LPi cell types cover the lobula plate with different scales, from ∼10% to nearly the entire field of view of each eye. Here we report a third instance of motion opponency, whereby the aggregate responses of the complete ipsilateral population of LLPC1 neurons appears to inhibit the complete population of LPC1 neurons through a single GABAergic interneuron in the central brain. This large-field inhibition occurs at the scale of the complete field of view of each eye (Fig. 7F). And finally, our consistent observation of contralateral motion influencing the encoding of ipsilateral motion signals in both LPC1 and LLPC1 demonstrates yet another inhibitory pathway, that combines visual information across the field of view of both eyes. The sharp selectivity of LPC1 and LLPC1 to translational stimuli therefore emerges from a cascaded arrangement of inhibition by non-preferred motion signals at increasing scales of vision.

## ACKNOWLEDGMENTS

The screen of Generation 1 GAL4 lines was carried out as part of the Fly Olympiad Project Team at Janelia with the assistance of James McMahon, Amanda Cavallaro, and others in Janelia’s Fly Core team, and also Nan Chen, Darlene Evich, Yukiko Ho, Mary Phillips, and Sharon Low. The GAL4 line screen benefitted from the guidance of the steering committee (including Julie Simpson and Gwyneth Card), early access to the FlyLight Project Team’s data, and Arnim Jenett and Yoshi Aso’s selection of GAL4 driver lines. We thank the Janelia FlyLight Project Team for help imaging driver lines. The Fly Vision Box was developed and supported with contributions from Steve Sawtelle, Sam Watkins, Magnus Karlsson, and software pipeline and analysis contributions from Allen Lee, Lowell Umayam, Frank Midgley, Don Olbris, Rob Svirskas, and Will Dickson. We thank Lihi Zelnik-Manor, Tim Lebetsky, Pietro Perona, and David Anderson, for permission to use and update fly tracking code that the Fly Vision Box video analysis is based on. We thank Emily Nielson for the schematic of the Fly Vision Box apparatus (Fig. 1A) and summary data visualizations. We are also grateful to members of the Reiser Lab, especially Kit Longden for support during this project and comments on the manuscript. This work is funded by the Howard Hughes Medical Institute through its support of the Janelia Research Campus. This article is subject to HHMI’s Open Access to Publications policy. HHMI lab heads have previously granted a nonexclusive CC BY 4.0 license to the public and a sublicensable license to HHMI in their research articles. Pursuant to those licenses, the author-accepted manuscript of this article can be made freely available under a CC BY 4.0 license immediately upon publication.

## AUTHOR CONTRIBUTIONS

Conceptualization, M.D.I., A.N., G.M.R., K.B., and M.B.R.

Methodology, M.D.I., A.N., G.K.L., T.T., A.W.E., W.K., K.B., and M.B.R.

Software: M.D.I., G.K.L., W.J.R., K.B., and M.B.R.

Validation: W.J.R., A.W.E., and W.K.

Formal Analysis: M.D.I., J.L.M.E., K.B., and M.B.R.

Investigation: M.D.I., J.L.M.E, A.N., W.J.R., and A.W.E.

Resources: A.N., E.M.R., and W.K.

Data Curation: W.J.R. and A.W.E.

Writing – Original Draft: M.D.I., A.N., and M.B.R.

Writing – Review & Editing: M.D.I., A.N., E.M.R., and M.B.R.

Supervision: W.K., G.M.R., and M.B.R.

Funding Acquisition: G.M.R., K.B., and M.B.R.

---

**Fig. S1, related to Fig. 1.**
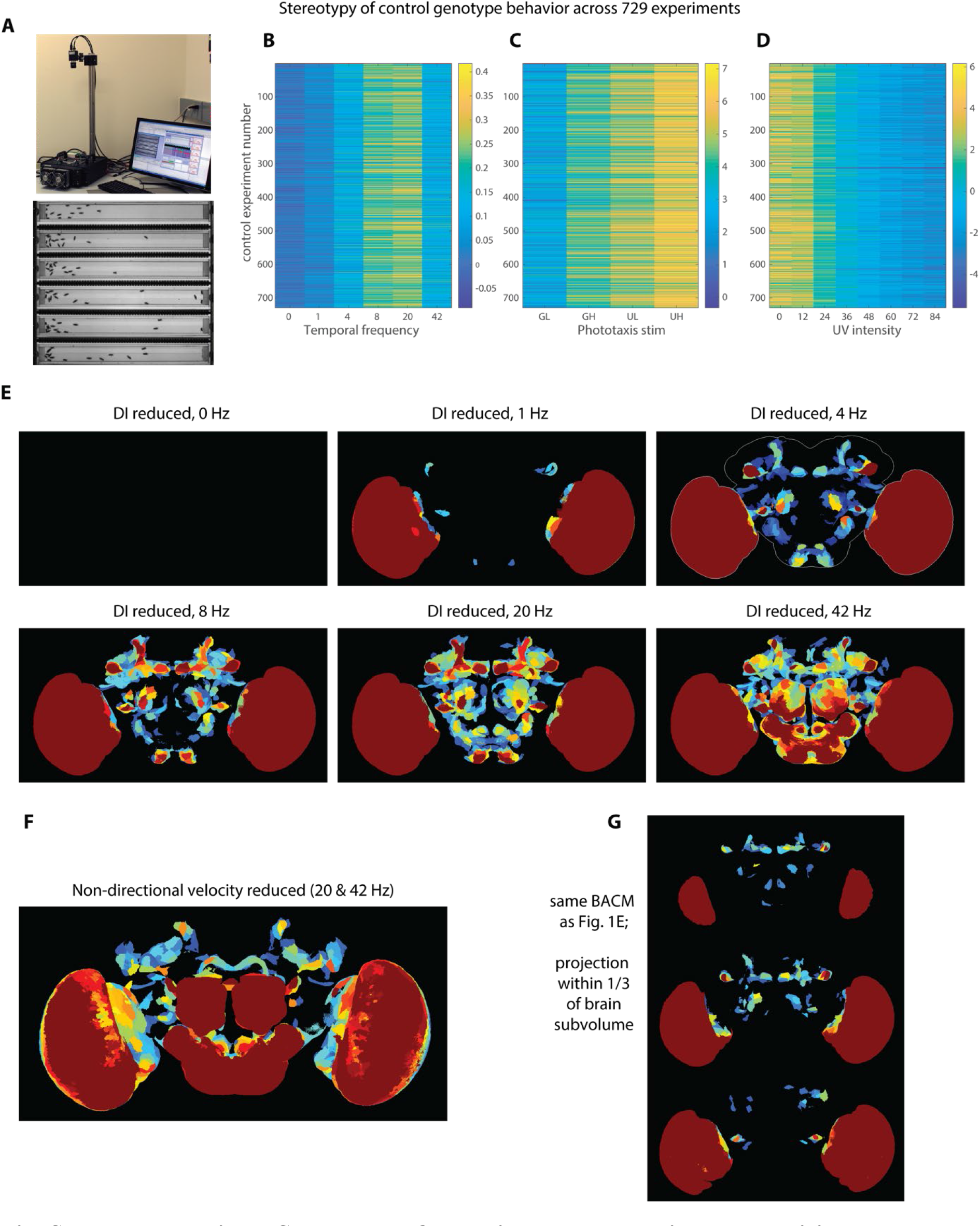
Stereotypy of behavioral responses in the Fly Vision Box and additional Behavior Anatomy Correlation Maps. (A) A photograph of the Fly Vision Box in the lab, showing the software interface, above a single frame view of a typical experiment. The tubes are 120 mm long and 10 mm wide. To further demonstrate the replicability of the fly behaviors we measure during our screen, the mean behaviors of >700 experiments of control flies are shown using a color-map to represent the behavioral metrics for different experimental modules of our screening protocol (see methods). (B) Responses to visual motion (as in Fig. 1 B, C) represented as the Direction Index (DI). (C) The phototaxis responses while flies walked towards UV or green LED with the indicated intensity level (GL – Green Low, GH – Green High, UL – UV low, UH – UV high) at the end of the tubes. The behavior is quantified as the Direction Index integrated over each trial (see methods). (D) The behavior of flies during a spectral preference task, similar to phototaxis, but where at one end a UV LED was illuminated with the indicated (relative) intensity, while the other end was illuminated with a green LED at a fixed intensity. The data are again represented as integrated Direction Indices. For the lower UV levels, flies walked towards the green LED, but walk towards the UV LED at higher UV levels. Behavior Anatomy Correlation Maps^46^ can be computed for any behavioral metric. In (E) the BACM for a reduction in the visual motion following behavior, computed as a Direction Index, for each temporal frequency, corresponding to each point along the tuning curve of Fig. 1C, Fig. S1B, is shown. The protocol contains a trial in which the grating pattern is shown but does not move. Flies could not generate a directional response to a non-moving pattern, and so the empty BACM corresponding to this ‘0 Hz’ condition serves as a sanity check for the analysis method. All BACMs for reduced directional following in the conditions with visual motion show correlations with expression in the optic lobes, while increasing speeds appear to be affected by contributions from broader expression throughout the central brain. The conditions of 4Hz, 8 Hz, 20 Hz, and 42 Hz all show expression in the PLP glomerulus, which became the focus of this study. The BACM in (F) corresponds to behavioral responses to visual motion that do not account for the direction of walking, as the DI metrics do. We note this BACM shows some similar, but also notably different units of expression the those in E. (G) shows the same BACM, for visual motion following as in Fig. 1E-lower, but shown here as 3 maximum intensity projections in each subvolume, roughly 1/3 the depth, of the fly brain. The color map in all BACMs is the same as in Fig. 1, logarithmically encoding the range of p-values from 0.05 to 0.0001.

**Fig. S2, related to Fig. 2.**
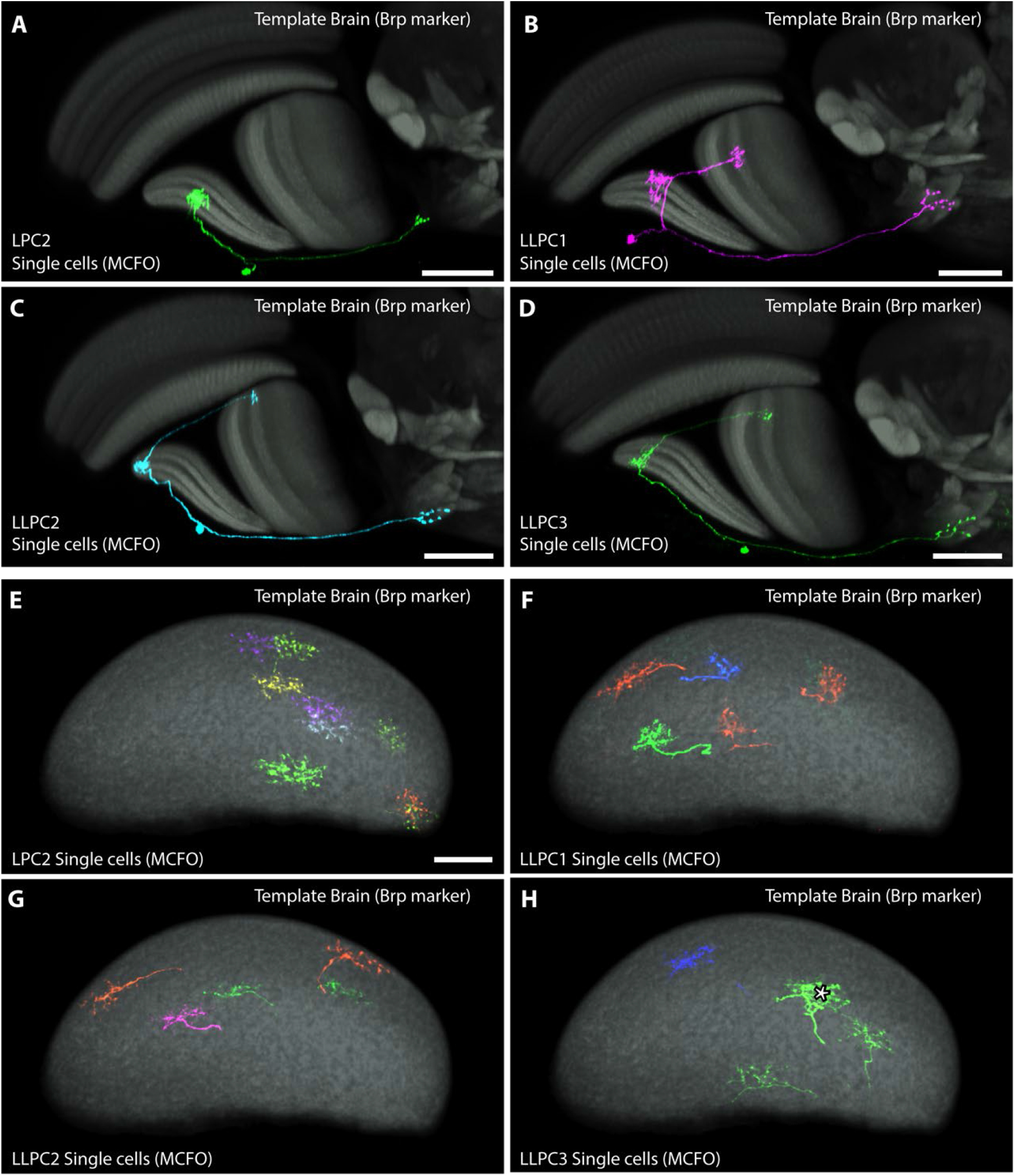
Single cell anatomy of the small-field Lobula Plate projection neurons. (A-D) Single cell anatomy of LPC2 (A), LLPC1(B), LLPC2(C) and LLPC3(D) cells. MCFO-labeled neurons of the indicated cell types displayed as in Fig. 2F. The field of view and projected volume are slightly different for each cell type to accommodate difference in the position of the labelled cells within the lobula plate. Scale bar, 30 µm. (E-H) En face views of the lobula plate with individual LPC2 (E), LLPC1(F), LLPC2(G) and LLPC3(H) cells. Asterisk in (H) marks two overlapping arbors from different cells. Images are composites of MCFO labeled single cells from multiple optic lobes displayed as in Fig. 2J. Scale bar in A-D is 30 µm, and 20 µm in E.

**Fig. S3 related to Figs. 2, 3.**
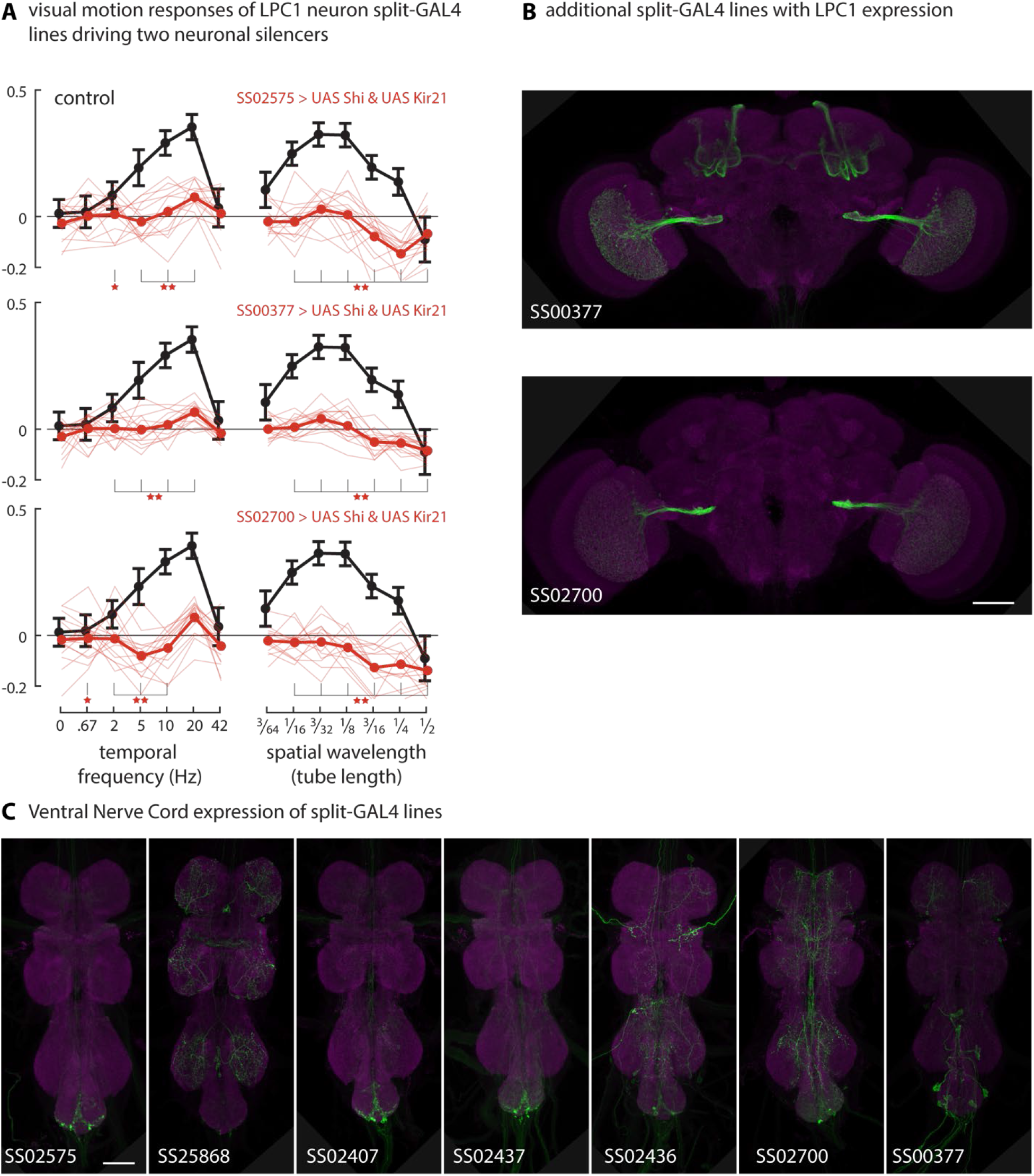
Fly Vision Box motion responses for additional LPC1 driver lines, and extended expression patterns for split-GAL4 lines. (A) Summary data for 3 additional split-GAL4 lines targeting expression in the LPC1 neurons. In these experiments, the neurons were ‘doubly silenced’ using both Shibire^ts1^ and Kir2.1, expected to interfere with distinct aspects (synaptic transmission and membrane excitability) of neuronal function. These data are plotted following the conventions of Fig. 3B: control genotype mean ± s.d. in black, and for the 3 experimental genotypes, the individual tube mean Direction Index is shown as thin lines, and the means across tubes as thicker lines (15-18 tubes from 3 experiments for all experimental and control genotypes). Statistical significance between experimental and control genotypes determined using Mann-Whitney U-test controlled for False Discovery Rate: 1 star = p < 0.05; 2 stars = p < 0.01. (B) Expression patterns of the two additional LPC1 driver lines (displayed as in Fig. 3A). Scale bar, 50 µm. (C) Ventral Nerve Cord expression of the driver lines in Figs. 3 and S3. Scale bar, 50 µm. Genotypes used in this Figure detailed in Table 1.

**Fig. S4, related to Fig. 4.**
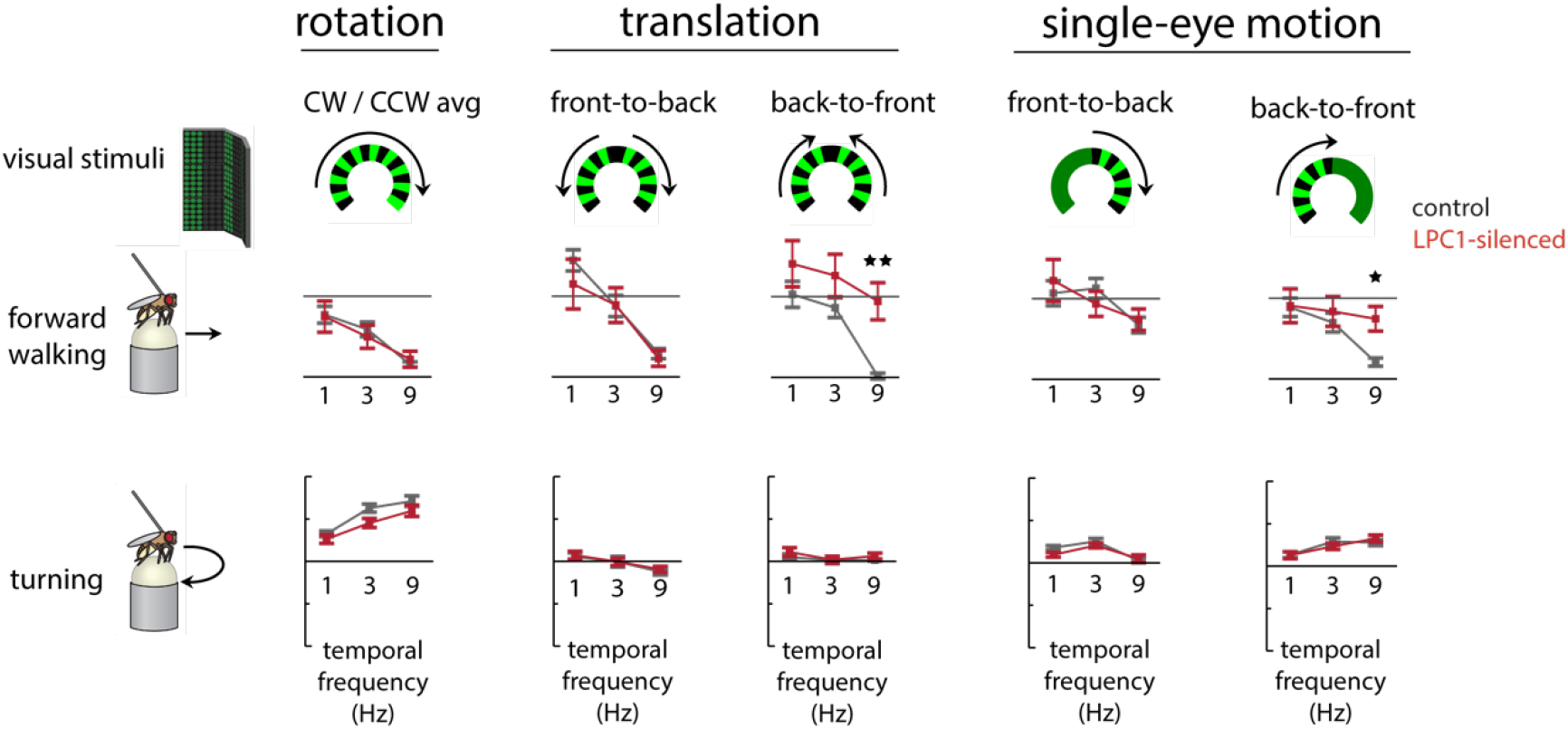
Visual motion responses of flies with LPC1 neurons silenced. A) Top: Grating patterns presented to tethered flies using a cylindrical LED arena (stimuli schematized with top-down view of arena). Middle: The forward component of the walking responses of tethered flies (n=20 LPC1-silenced flies in red; n=20 genetic control flies in black) to presentations of rotational and translational motion stimuli (at 1 Hz, 3 Hz, and 9 Hz temporal frequency, presented for 2 s), displayed as mean ± SEM. Bottom: Mean turning component of the walking behavior of flies, from the same data set as above. LPC1-silenced flies walked significantly more (or slowed down less) than control flies only in response to 9 Hz full-field and 9 Hz single-eye back-to-front motion (statistical significance determined using Mann-Whitney U-test controlled for False Discovery Rate: 1 star = p<0.05; 2 stars = p<0.01).

**Fig. S5, related to Fig. 5.**
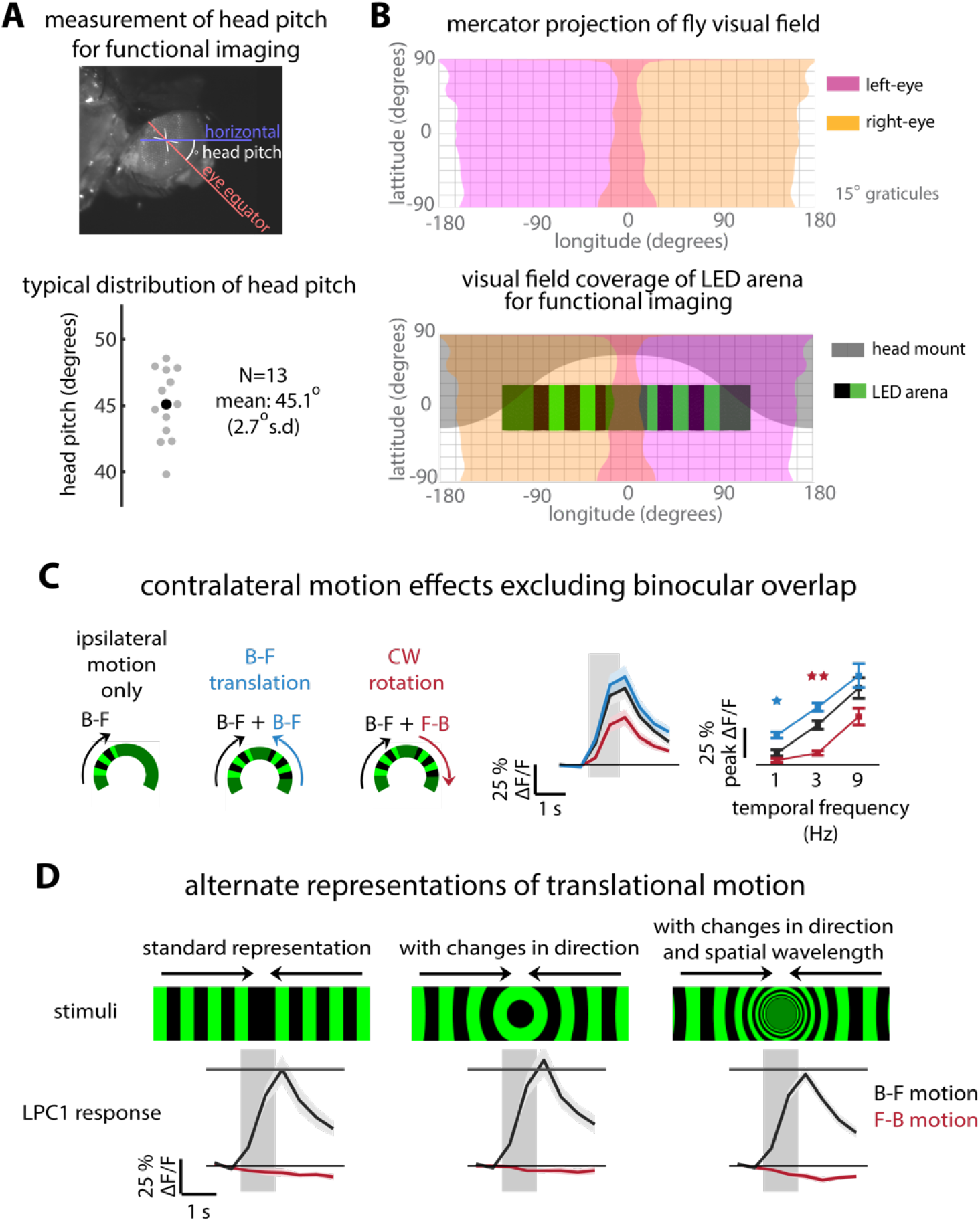
Visual stimulus design considerations for measuring translational motion responses in LPC1 neurons. (A) Measurement of head angle of flies mounted for functional imaging experiments. The fly head is typically pitched 40-50° downward to allow access to the posterior side of the brain. (B) Approximate mapping of the fly’s field of view during the imaging experiments, used to inform stimulus design (based on prior work^67, 68^). For monocular stimuli, designed to be visible by only one eye, the motion patterns are designed to start 20° from the midline of the fly’s field of view to avoid the region of binocular overlap. Small-field stimuli (shown in Figs. 5B, 6B, and 7E) were designed to stop at 90° from the midline (directly to the side) to avoid occlusions by the head mount, which could cause some confounding motion cues. (C) The left LPC1 population responses are significantly modulated by the direction of motion presented to the contralateral eye, as in Fig. 5E, but here for stimuli that exclude the region of binocular overlap (n=10 flies). (D) Forward/backward translational stimuli can be rendered with higher fidelity on our cylindrical display by including changes in direction and spatial wavelength. The calcium responses of LPC1 neurons (plotted as mean ± SEM; n=6 flies) is not significantly altered by different types of back-to-front translation stimuli. Based on the set of stimuli we tested, the horizontal component of the translational motion and the combination of left/right directions on each eye appear to be the primary determinants of LPC1 population responses.

**Fig. S6 related to Fig. 6.**
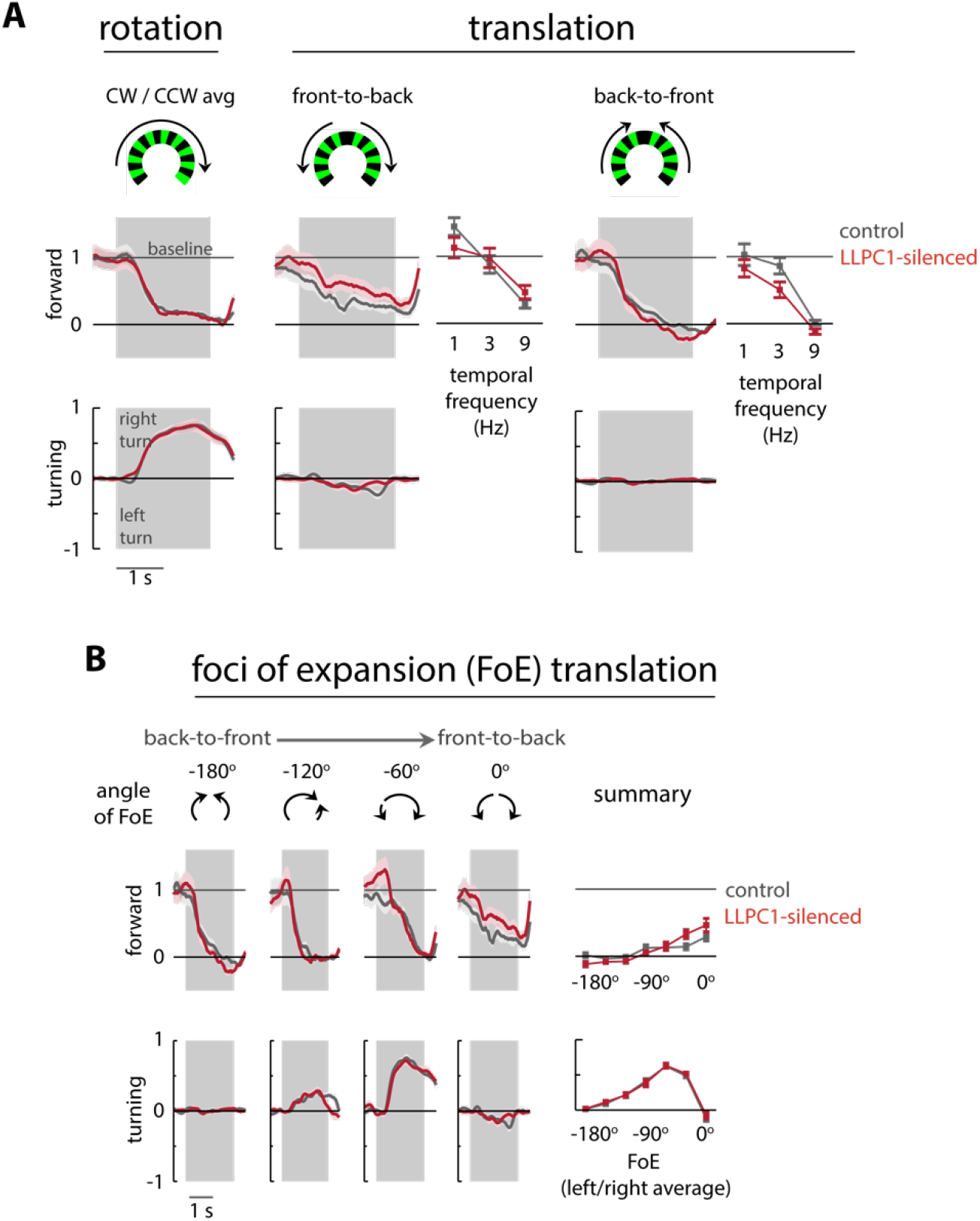
Silencing LLPC1 neurons did not affect visual responses of tethered flies walking on an air-supported ball. (A, B) Changes in forward and turning components of walking responses to rotational and translation visual motion stimuli for control flies and flies with LLPC1 neurons silenced by expressing Shi^ts1^. The experiments were identical to those conducted on LPC1-silenced flies (Fig. 4B, C). For this n=17 flies data set, we find no statistically significant effects on either the forward or turning components of the walking behavior in response to any of the presented stimuli.

**Fig. S7, related to Fig. 6.**
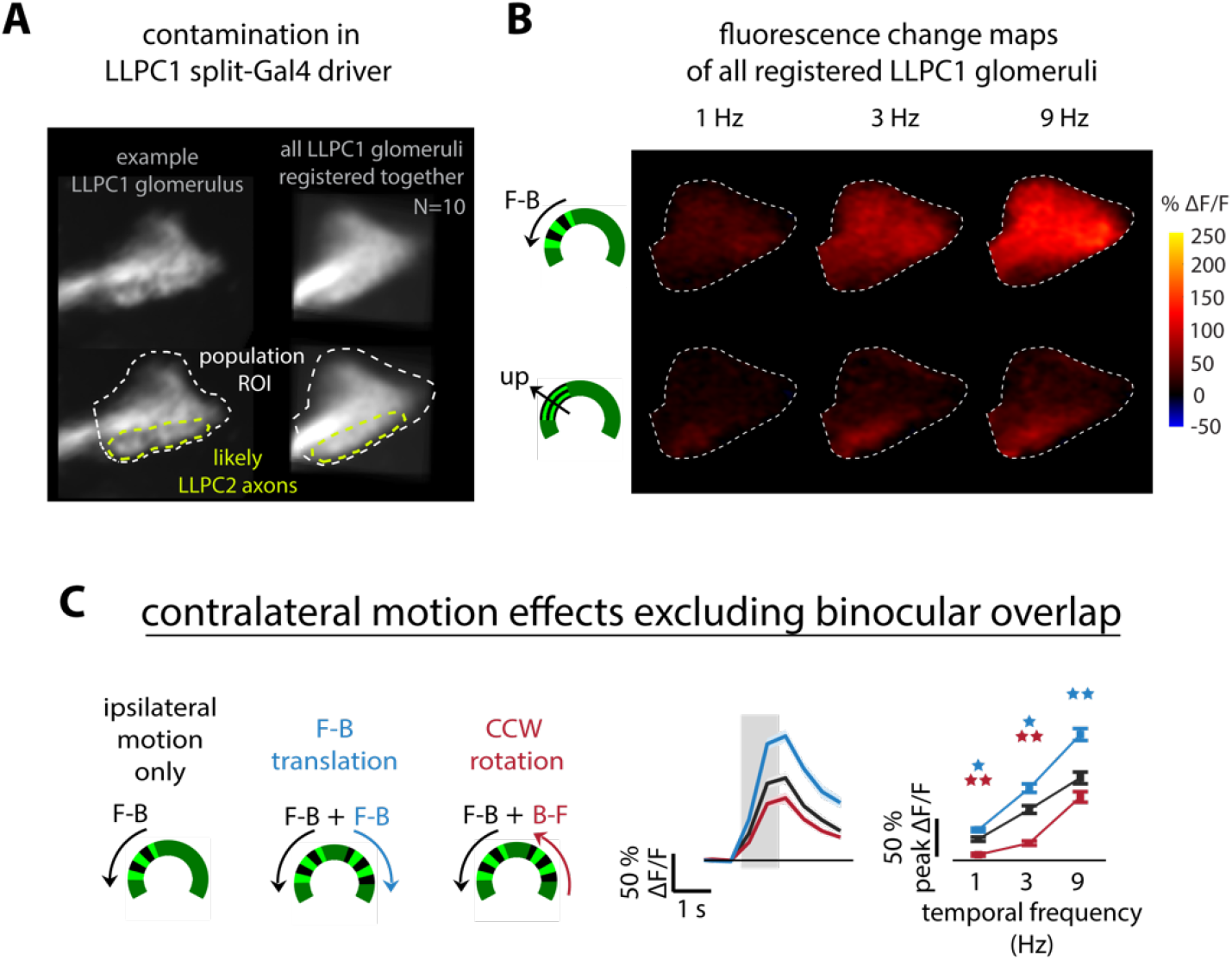
Expanded results from LLPC1 imaging experiments. (A) Functional imaging of LLPC1 with genetically targeted expression of GCaMP6m is likely contaminated by the axons of LLPC2 neurons present in the inferior margin of the LLPC1 glomerulus. (B) The superior side of the LLPC1 glomerulus is strongly activated by front-to-back motion, as expected from the inputs LLPC1 dendrites receive in Lobula Plate layer 1 (Fig. 2). In this driver line (see Table 1), the inferior margin of the LLPC1 glomerulus is consistently most selective to upward motion (analysis summarizes measurements from the registered glomeruli of n=10 flies). The LLPC1 split-GAL4 driver line likely contains some LLPC2 neurons, explaining this mixed selectivity, but with spatially separated responses indicating distinct populations of neurons. (C) The left LLPC1 population responses are significantly modulated by the direction of motion presented to the contralateral eye, as in Fig. 6E, but here for stimuli that exclude the region of binocular overlap, similar to LPC1 (Fig. S5C) but for the opposite directions of motion (n=10 flies).

**Fig. S8, related to Fig. 7.**
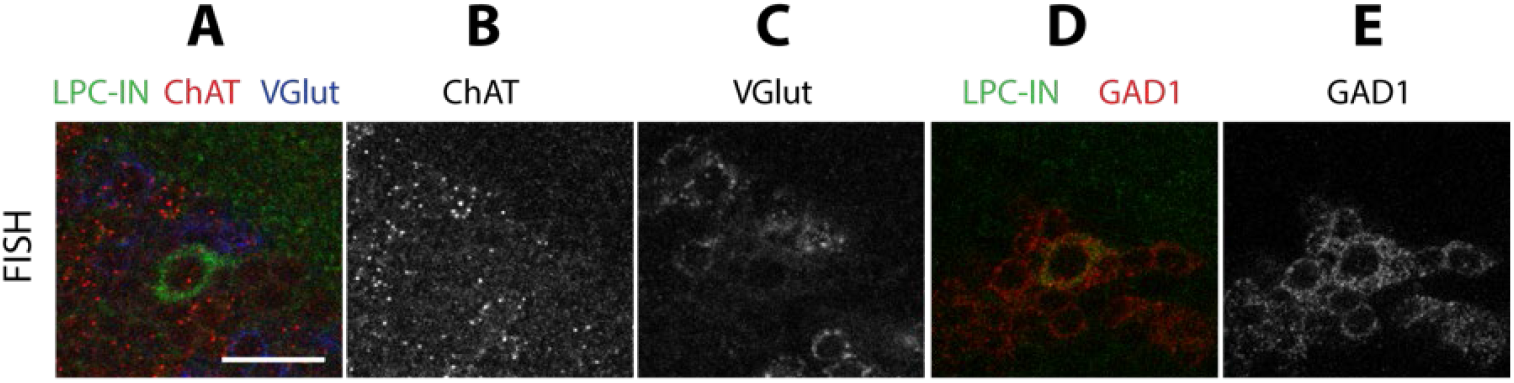
FISH labelling supports a GABAergic phenotype of the LPC-IN neuron. FISH (florescence in situ hybridization) labelling^69^ reveals the likely neurotransmitter phenotype of the LPC-IN cell. The LPC-IN cell (green) shows above background FISH signal with probes for GAD1, but not for ChAT and VGlut, suggesting that this cell type is GABAergic. Scale bar, 10 µm.

## METHODS

### Flies

Flies were reared under standard conditions (60% humidity, 16h light / 8h dark, cornmeal agar diet) at 22° C for Fly Vision Box, tethered walking, and functional imaging experiments. Flies expressing CsChrimson were reared on a cornmeal agar diet supplemented with 0.4 mM all-trans-retinal. Fly Vision Box experiments were performed on male flies 4-5 days post-eclosion, imaging experiments on female flies 1-4 days post-eclosion, and tethered behavioral experiments on female flies 3-5 days post-eclosion. Cell type specific expression of indicators, silencers, and activators was achieved using the GAL4/UAS, LexA/LexAop, and Split-GAL4/UAS expression control systems^44, 52, 53^. We used previously described split-Gal4 driver lines^54^ or constructed new drivers from existing hemidriver lines^70, 71^. Expression of CsChrimson^58^ in bilateral or unilateral (or empty) LPC1 populations was achieved using the FLP/FRT expression control system^72^, following a previously introduced method^47^. Tables 1 and S1 detail the complete list of the fly genotypes used in this study.

### Anatomical analyses

For imaging of split-GAL4 expression patterns, we used MCFO, a method for stochastic labeling of neurons in multiple colors, to visualize the anatomy of individual cells and targeted expression of membrane and presynaptic markers to reveal cell populations^47, 65^. Sample preparation and imaging were carried out by the Janelia FlyLight Project Team. Protocols were as in previous studies^47, 65^ and are also available online (https://www.janelia.org/project-team/flylight/protocols). FISH detection of transcripts of GAD1, VGlut and ChaT used a recently described method^69^. Details of fly genotypes are provided in Table 1.

Images for anatomy figures were processed and displayed using Fiji^73^ and FluoRender^74^. A brief summary of image processing for each figure panel is included in Table 1. To generate composites of multiple confocal images, image stacks were first registered to a template brain (JRC2018U)^75^. In a few cases, stacks were edited to exclude signal from other cells or background signals that would otherwise obscure relevant details in the views shown. Such editing was not applied to the images of overall expression patterns (Figs. 3 and S3). Original image stacks will be made available online at https://splitgal4.janelia.org/cgi-bin/splitgal4.cgi.

### Preparation for Fly Vision Box experiments

Male flies of each genotype were cold-anesthetized and sorted into groups of 10-15 flies, then placed on “starvation media” (10-15 ml of 1% agar in water, in a standard fly vial), 1-4 hours prior to the start of experiments. Each group of flies was loaded into a custom-made clear acrylic corridor tube (interior dimensions: 3 mm ×10 mm × 120 mm, H × W × L) using a custom-made transfer-funnel.

### Preparation for tethered walking experiments

Flies were prepared for walking behavior experiments as previously detailed^13^ and briefly summarized here. Wings of cold-anesthetized flies were immobilized with UV-curing glue (KOA 300-1; Kemxert) at least 12 h prior to the experiments. On the day of experiments, each fly was glued to a tungsten rod (catalog #71600; A-M Systems) by the thorax and placed into an incubator (model PH09-DM, Thermoelectric Series; Darwin Chambers Company) set to 32°C and 60% humidity for at least 25 min before the start of behavioral experiments. We recently detailed this method and lower-cost alternatives^76^.

### Preparation for calcium imaging

Flies were prepared for 2-photon calcium imaging experiments using a method based on previously described preparation^55^ with several modifications as summarized here. Cold-anesthetized flies were tethered to a fine tungsten wire using UV-curing glue (Loctite 3972; Henkel). The two most anterior legs (T1) were severed and sealed with glue to prevent the fly from grooming and obstructing its visual field. The tethered fly was positioned up to the opening of a custom-machined PEEK plastic conical mount with its head pitched downward to allow access to the back of the head for dissection and imaging while obstructing as little of the fly’s visual field as possible^41^. The fly was glued to the mount at the rim of the head and the upper thorax, sealing most of the mount’s opening. The proboscis was glued to the severed legs to further immobilize the head. The back surface of the fly’s head was bathed in saline^55^ and a hole in the cuticle was cut to expose the posterior lateral protocerebrum (PLP) region of the brain using a fine tungsten needle and holder (Fine Science Tools #10130-05, # 26018-17). Muscles 1 and 16^77^ were severed to reduce motion of the brain within the head capsule and excess fat was removed from the surface of the brain to improve optical access. The complete procedure is shown in Movie S1.

### Optogenetics stimulation and experimental procedure

For optogenetic activation of CsChrimson (Fig. 4D), a 660 nm fiber-coupled LED (Thorlabs Deep Red M660F1) controlled by an LED driver (Thorlabs LEDD1B), set to ∼0.7 A, was directed approximately 1 inch from the tethered fly with a light guide (400 µm, 0.39 NA SMA to ceramic ferrule patch cable, Thorlabs FT400EMT). The LED was triggered using a signal, whose timing was controlled by analog output signals of our “G3” display controller^56^. After each optogenetic experiment, the fly brain was dissected and imaged with an epifluorescence microscope to record the location of CsChrimson expression (i.e., left optic lobe, right optic lobe, both, or neither).

### Visual stimuli

For the fly-on-ball experiments of Figs. 4, S4, S6, visual stimuli were presented to tethered flies using a 3^rd^ generation, or “G3” cylindrical LED arena (manufactured by IO Rodeo, based on the design detailed at: https://reiserlab.github.io/Modular-LED-Display/G3/) constructed from 36, 8×8 pixel LED modules containing green LEDs (UltraPure Green LED, IO Rodeo, 525 nm peak). The cylindrical arena covered 270° across the azimuth of the visual field and ∼120° in elevation. The maximal angular distance between pixels, determined from the flies’ position is 3.75°. This system is slightly updated from the published, earlier version^56^.

For calcium imaging experiments, visual stimuli were presented to tethered flies using a 4^th^ generation “G4” cylindrical arena constructed from 36 16x16 pixel LED panel containing green (570 nm peak) LEDs^41^. The LED arena was pitched downward to match the head angle of the fly determined under the dissection microscope so that the LED arena was aligned to the equator of each the fly’s eyes (Fig. S5A,B). To further separate the yellow-green LED emission spectra from GCaMP6m fluorescence, a long-wavelength passing gel filter (Deep Golden Amber, LEE Filters #135) was laid on top of the LEDs along with a diffusing filter (Full White Diffusion filter, LEE filters #216) to prevent reflections from the gel filter.

Patterns displayed on either LED arena were generated using a set of MATLAB scripts for parameterizing and visualizing moving grating patterns for this display system. These tools have since been integrated into a GUI (described here^41^ and available here: https://reiserlab.github.io/Modular-LED-Display/G4/). All experimental stimuli consisted of 100% contrast square-wave gratings with a 30° spatial frequency. Brightness of stimuli was modulated on a 0-15 (4-bit) range, with values of 15 and 0 typically used for bright and dark bars in grating stimuli and a value of 7 for background and intermediate brightness frames. For optomotor experiments with CsChrimson flies (Fig 4D), the display brightness was lowered to reduce potential unintended optogenetic activation from the display, whose green LEDs emit some longer wavelength light (brightness values of 2/1/0 were used for bright bar/background/dark bars). For tethered walking experiments, each visual stimulus trial began with 2 seconds with the display set to a uniform, intermediate brightness (overall brightness does not substantially change when the gratings appear), followed by 2 seconds of the moving grating stimulus, followed by another 2 seconds of the intermediate brightness. For optogenetic activation experiments, no grating pattern was displayed, but the uniform, intermediate brightness pattern was displayed for 6 seconds with the 660nm LED turning on after 2 seconds and remaining illuminated for 1 second. For calcium imaging experiments, a single trial consisted of 2 seconds of uniform, intermediate intensity, followed by 1 second of moving gratings, followed by 6 seconds of the uniform, intermediate display. The increased time in between stimulus trials was added to allow GCaMP6m fluorescence signal to return to baseline levels. Experiments were organized using a randomized block trial structure. Within each block, the complete set of visual stimuli were presented in a randomized order, and multiple blocks were presented such that each stimulus was shown at least 3 times. For imaging experiments, every trial was repeated 3 times; for most behavioral experiments, trials were repeated 4 times; and for optogenetic activation experiments (1 s stimulation by the 660 nm LED with the uniform visual stimulus), trials were repeated 15 times.

### Fly Vision Box experiment

The Fly Vision Box was developed at the Janelia Research Campus as a high-throughput assay integrating several tests of visually guided behavior for flies in tubes, including phototaxis, visual motion following, and color preference. The transparent tubes were custom made from acrylic (interior dimensions: 10 mm × 3 mm × 120 mm, W × H × L), to allow flies a corridor to run up/down, while preventing flies from walking over each other, simplifying both the behavior and our ability to track it. Each tube is capped on both end with a custom machined, transparent acrylic plug that solidly seals each corridor. Each endcap also houses a 0.125” cube magnet, glued into a machined slot, for alignment of each tube into the box’s lid. 5 boxes in total were built, 4 of these were used to conduct experiments for the GAL4 line screen summarized in Figure 1 and Table S1.

The apparatus is a temperature-controlled box that holds 6 tubes. Temperature control was implemented via 2 Peltier elements mounted on the side of the box, with fans mounted on either side for air exchange. On either side of every tube is a row of 64 green (572 nm peak), surface mount LEDs, mounted on a custom designed Printed Circuit Board (PCB). The PCB houses an Atmega168 microcontroller and is essentially an elongated G3 LED panel^56^ (but supporting 2 LEDs per pixel, mirrored on each side of the board), scanning the LEDs and being controlled using the same code and protocol. A single green (564 nm peak) and a single UV (350 nm peak) LED are mounted on a small PCB at each end of each corridor. These LEDs are intensity controlled for phototaxis and color preference experiments by another Atmega microcontroller (ATmega644, Atmel). The tubes are suspended from the top-plate that seals the box (and aligned by magnets onto posts). 4 small vibrating ‘pager’ type motors are mounted onto the sides of the top plate to provide a mechanical startle for the walking flies. A camera (Basler A602f) mounted above the box records fly movements at 25 frames/s. The camera is fitted with an Infrared-passing filter and the tubes are suspended above an Infrared backlight (Advanced Illumination, 880 nm LEDs). The camera records videos that are stored for off-line tracking and analysis.

The Fly Vision Box is controlled via software that provides deterministic timing of the experimental protocol. After many iterations to optimize the reliability of the measured behavioral responses of control flies, the experimental protocol (v3.1) that was run for the Gen1 GAL4 line screen consisted of 5 sequences, run twice, once at 24°C and a second time at an elevated temperate of 34°C:

1. 180 s acclimation period, during which the flies are in the dark. The last 2 minutes of the trial are recorded.
2. 180 s test of mechanical startle responses. Every 30s a 100 ms vibration is delivered by the 4 pager motors. 6 repetitions total (in subsequent trials, the same 100 ms vibration is paired with the start of each condition).
3. 240 s test of visual motion responses. Every 10 s, a vibration is delivered by the pager motors, and a visual motion stimulus is presented at one of 6 speeds (0, 1, 4, 8, 20 and 42 Hz). The pattern is composed of 2 LED pixels on, and then 2 pixels off, running the length of the tube. The update rate is calculated such that these temporal frequencies are displayed (pattern shifts by one pixel every 0, 250, 63, 31, 13, and 6 ms respectively). The motion is directed towards one end of the tubes, then towards the other end in the subsequent 10 second trial. The complete protocol includes 4 repetitions of each speed. Control behavior summarized in Figure S1B.
4. 120 s test of phototaxis behavior. Every 15 s, a vibration is delivered by the pager motors, and a single LED is illuminated at one end of the tube, before the same color LED is illuminated on the opposite side of the tube. There are 4 conditions (GL: Green Low, 25; GH: Green High, 120; UL: UV low, 36; UH: UV high, 200). The complete protocol includes 4 repetitions of each condition. Control behavior summarized in Figure S1C.
5. 240 s test of color/spectral preference. Every 15 s, a vibration of the pager motors is accompanied by a condition similar to phototaxis, but where at one end a UV LED was illuminated with a different (relative) intensity (0, 12, 24, 36, 48, 60, 72, 84), while the other end was illuminated with a green LED at a fixed intensity (120). Control behavior summarized in Figure S1D.

After this screen was carried out, we expanded the protocol to measure these behaviors more thoroughly. During this process, we found that we could not reliably measure visual responses to lower contrast visual motion stimuli. Our understanding is that flies experience these optomotor patterns as high-brightness but already low-contrast stimuli due to internal reflections and the opposing rows of LEDs illuminating each other. To overcome this limitation, the updated protocol (v5.27 and v5.34, identical but with the order of sequences changed) was carried out with one side of each tube’s LEDs covered with black laboratory tape. The enhanced protocol consisted of 8 sequences, carried out while the box is heated to 34°C (no 24°C experiments, and the entire duration is slightly variable since the experiment doesn’t begin until the box has achieved the elevated temperature):

1. A “startle” response test identical to sequence 2 from the protocol above.
2. 280 s visual motion response test leading to a temporal frequency tuning curve, as above, except that the moving pattern consisted of 4 LEDs ON followed by 4 LEDs OFF (repeated down the length of the tube), scrolling at 7 temporal frequencies (0, 0.67, 2, 5, 10, 21, and 42 Hz). Each stimulus was repeated 2 times in both directions for 4 total repetitions.
3. 200 s visual motion series testing different contrast levels over a mean intermediate luminance. The pattern is 4 brighter pixels / 4 darker pixels repeating down the length of the tube, moving at 5 Hz temporal frequency. 4-bit brightness intensity control allows possible brightness values of 0-15. 5 contrast levels were tested: 0.07 (8/7), 0.2 (9/6), 0.5 (11/4), 0.7 (13/2), and 1 (15/0). Each stimulus was repeated 2 times in both directions for 4 total repetitions.
4. 200 s visual motion series testing different contrast levels, as above, but where the 5 stimuli were chosen to maintain the brightest level: 0.1 (15/12), 0.3 (15/9), 0.4 (15/6), 0.7 (15/3), and 1 (15/0). Each stimulus was repeated 2 times in both directions for 4 total repetitions.
5. 280 s visual motion response test leading to a spatial wavelength tuning curve. 7 different patterns composed of the following repeating sequences of on/off pixels were moved: (1/2, 2/2, 3/3, 4/4, 6/6, 8/8, and 16/16, corresponding to 3/64, 1/16, 3/32, 1/8, 3/16, 1/4, and 1/2 the length of the tube), and the speed of movement was adjusted to approximate 10 Hz temporal frequency: (32, 24, 16, 12, 8, 6, 3 ms per step, respectively). Each stimulus was repeated 2 times in both directions for 4 total repetitions.
6. 120 s test of phototaxis behavior, same as seq 4 in v3.1, but here (GL: Green Low, 20; GH: Green High, 120; UL: UV low, 15; UH: UV high, 200).
7. 440 s test of spectral preference, similar to seq. 5 in v3.1, but here the trials are 10 s each, and 11 different combinations are shown, where the UV LED is held at a constant intensity. On one side, the UV LED is set to 10 while the green on the other end is set to 11 values (0, 3, 6, 10, 15, 20, 30, 50, 75, 100, and 200). Each combination was repeated 2 times in both directions for 4 total repetitions.
8. A 440 s test of spectral preference, as in (7) above, but here the green LED is held at a constant intensity of 100 while the opposite—side UV LED is set to 11 values (0, 5, 7, 10, 15, 25, 40, 55, 75, 100, and 200).

### Fly Vision Box analysis

After each experiment was completed, the movies were analyzed offline. Extensive metadata and protocol information (genotype, sex, DOB, age, time of day, rearing conditions, etc.) were stored along with the experimental data. To minimize storage space and accelerate subsequent analysis, the videos of all 6 tubes in each box were split into videos of individual tubes, and then further compressed into static-background SBFMF files^78^. Fly positions were detected in each video frame using background subtraction, and fly movements were tracked across frames using software developed by Lihi Zelnik-Manor and Pietro Perona and used in prior work^79^. The tracking and analysis code has been updated and maintained by Janelia Scientific Computing and is available here: https://github.com/JaneliaSciComp/box. Since the tracking software did not preserve the identity of individual flies, the most reliable metrics of directed walking behavior were based on the (directional, median) walking velocity of all flies in each tube. We further found that a direction index was an even more robust measure of behavior, that was less confounded by experiment-to-experiment and genotype-to-genotype differences in walking speeds. The direction index is defined as 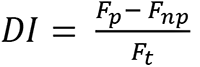, where *F_p_* is the number of flies moving toward the preferred direction, *F_np_* is the number of flies moving in the nonpreferred direction, and *F_t_* is the total number of flies detected in the corridor. The preferred and nonpreferred directions are defined for each stimulus (opposite to the direction of motion for the visual motion task, and towards the illuminated LED in the phototaxis behaviors). The DI is computed for each tube, frame-by-frame, and then across trials.

The data in Figure 1 is based on the motion responses in sequence 3 of protocol v3.1, from the second half of the experiment, when the box is held at an elevated (34°C) temperature. The temporal frequency data and the spatial wavelength data in Figure 3 and Figure S3, are from the responses in sequence 3 and 5 of protocol v5.27 (5 and 8 of v5.34).

The Behavior Anatomy Correlation Maps (Figure 1, and Figure S1) were produced using the methods and code previously described^46^. The figures were produced by the BABAM package (available: https://github.com/kristinbranson/BABAM), with settings similar to those used in prior work: 10,000 samples, False Discovery Rate (FDR) = 0.25, logarithmic colormap interpolation, min p-value = 0.0001, max p-value = 0.05 and are shown as maximum intensity projections of full-brain volumetric data. In Figure 1E, the top plot is based on the metric ‘MedVelDark_minus’ (median walking velocity in the dark) and the bottom BACM is based on ‘DI_Fast_minus_ANDNOT_ND_MedVel_Fast_minus’ – this compound result uses a logical operation on 2 metrics, a reduction (relative to control) in the direction index during the fast motion trials (20 and 42 Hz) and no reduction in non-directional walking during this same trials. This combination produces a slightly ‘cleaner’ map by eliminating regions that appear to affect walking during these fast motion trials, but not in a directional manner. The BACMs based on the individual metrics used in this analysis are shown in Figure S1, where the DI reduction BACM is shown for each walking speed (in E). At elevated speeds, the PLP ‘hotspot’ is clearly seen, but additional regions in the central brain also become prominent. However, the ‘recruitment’ of these regions may have more to do with high levels of locomotion from these agitated flies, and not with visual motion processing. The clearest evidence for this is the non-directional walking velocity reduction BACM shown in F, where a significant role for expression in e.g., the antenna lobes and the SEZ is found, and these are the overlapping regions that are reduced in the BACM of Fig. 1E, lower. All Behavior Anatomy Correlation Maps based on the visual motion behavior from sequence 3 of protocol v3.1. The screen data not shown in these figures is summarized in Table S1.

### Tethered walking equipment and analysis

The methods used for these behavioral experiments have been detailed elsewhere^13, 55^ and are summarized here. Tethered flies were positioned onto an air-supported foam ball using a 3-axis micromanipulator and two cameras (Firefly MV FFMV-03M2M and Basler 602f) that imaged the fly and ball from above and behind, for alignment. The foam ball was cut from FR-7120 material, measured 9 mm diameter and 129 mg, was supported by 340 SCCM airflow, regulated by a digital mass flow controller (Smart-Trak, Sierra Instruments). The complete setup, which included the visual LED arena^56^ and the treadmill system^55^ was housed within the incubator.

Behavioral experiments were conducted at 34°C and 60% humidity. The ball movement was tracked by two optical flow cameras (ADNS-6090; Avago Technologies) that measure image displacements. These measurements were acquired by a custom MATLAB (MathWorks, Natick, MA) program that runs the experiment, and these signals were transformed into 100 Hz measurements of forward, sideslip, and turning measurements (ball pitch, roll, and yaw), in arbitrary units. These measurements of displacement are scaled by 100/s to generate velocity signals (in arbitrary units per second).

Only turning and forward measurements were analyzed and described in this work. The time-series of turning and forward measurements in response to multiple repetitions of each visual stimulus were averaged together to produce a mean turning and forward response for each fly, and this was computed for all visual stimuli. For the mean time-series of turning, all flies of a single genotype were then normalized to the 98^th^ percentile turning measurement of all flies from that genotype (a more robust procedure than normalizing to peak), which scaled the values to range from approximately -1 to +1, where -1 represents an approximately maximum yaw left (counter-clockwise) turn and +1 indicates an approximately maximum yaw right (clockwise) turn. For the average time-series of forward walking, all flies of a single genotype were normalized to the mean forward measurement during the 0.5 seconds prior to all visual stimuli (at the end of the 2 seconds of intermediate, uniform brightness that preceded all experimental stimuli). This procedure scales the values such that +1 represents the baseline forward walking prior to the start of experimental visual stimuli and 0 represents no forward walking. When appropriate, behavior measurements from left/right symmetric stimuli are averaged together for each fly individually: For full-field clockwise and counterclockwise rotation (e.g. Figure 4B, left), forward walking responses to both stimuli are averaged, while turning responses are “flipped and averaged” (left turns from CCW stimuli sign flipped and averaged with right turns from CW stimuli). For single-eye stimuli (e.g. Figure 4B, right), forward walking from right-eye B-F stimuli is averaged with that of left-eye B-F stimuli, while left turns from right-eye B-F is flipped and averaged with left-eye B-F (single-eye F-B stimuli is processed similarly). For summary plots, a single value of the turning and forward responses was calculated from the time-series by averaging the mean time-series values during the 2^nd^ half of open-loop visual stimulus presentation, excluding the 1^st^ half of the open-loop section to ignore the visuo-motor delay period and transients. Both time-series and summary data are plotted as the mean of all flies of a single genotype +/-SEM.

### 2-photon imaging and analysis

Imaging the fluorescent calcium indicator signals from neurons expressing GCaMP in the posterior lateral protocerebrum (PLP) of the fly brain was conducted using a two-photon microscope (Prairie Ultima) with near-infrared excitation (930nm, Coherent Chameleon Ultra II) and a 40x objective (Nikon CFI APO 60XW). The excitation power never exceeded 25 mW at the sample. T-series of z-stacks (z-series) were taken at 128x128x5 pixel resolution (0.446 µm/pixel along x and y axes and 5 µm/pixel in z-axis, resulting in a 57x57x25 µm imaging volume) at a rate of 2.02 Hz, using galvo-galvo scanning in the x and y dimensions and piezo scanning in the z-dimension. For each experimental condition, 10 z-stacks were acquired, precisely synchronized to the visual stimulus. Acquisition was triggered to start 2 seconds before the experimental visual stimulus was displayed, using the on-board analog outputs of G4 display system: 4 z-stacks were recorded in the 2 seconds prior to the experimental stimulus, 2 recorded during the 1 second experimental stimulus, followed by another 4 after the experimental stimulus had ended.

The time-series of image volumes was converted into a time-series of 2D images by computing a mean z-projection. Collapsing the z-stacks minimizes the effect of any brain motion in the z-direction. Motion in the x and y directions was corrected using “imregister” in the MATLAB image processing toolbox: each image was first gaussian filtered (alpha=3 pixels) to improve registration of noisy images, then registered using the ‘multimodal’ metric and ‘rigid’ transformation. Using this motion-stabilized t-series, an ROI was hand-drawn over the cell/cell-population of interest. For LPC1 and LLPC1, an ROI was draw over the entire glomerulus population, excluding any additional axon tract that didn’t overlap with the glomerulus. For LPC-IN, an ROI was drawn over the area of most dense branching. The mean pixel value within the entire ROI was used as the population average fluorescence for each frame. Since each experimental condition resulted in 10 values for each experimental condition (4 values before the stimulus, 2 during, and 4 after), the fluorescence change (ΔF/F) was calculated by dividing all 10 fluorescence values by the baseline fluorescence – the mean of the first 4 ‘pre-stimulus’ values.

Peak ΔF/F was computed by finding the max value of frames 6-8 in each time-series; values 1-5 are ignored as they are the pre-stimulus fluorescence change values and frames 9-10 are ignored as the peak fluorescence change was never observed more than 2 seconds after the experimental stimulus began, due to the temporal dynamics of GCaMP6m and GCaMP6s. Both ΔF/F time-series and peak ΔF/F are plotted as mean of all flies +/-SEM.

To create fluorescence change maps (Fig. S7B), instead of calculating a time-series of ΔF/F using the mean value within an ROI as above, this calculation was performed on every pixel in the image independently and the fluorescence change value calculated for each pixel (for each set of images per stimulus) was then color-coded by the amount of fluorescence change (as indicated by the color-scale bar in Fig. S7B).

### Statistics

P-values were calculated using the Mann-Whitney U-test (“ranksum” function in MATLAB). We then controlled for the False Discovery Rate (FDR) associated with multiple comparisons using the (more conservative) Benjamini-Yekutieli procedure^80^ was used to correct for the false discovery rate (FDR) associated with multiple comparisons. For the split-GAL4 silencing experiments (Fig. 3), 70 comparisons (14 conditions × 5 genotypes) were made, and for the LPC1-double silencing experiments (Fig. S3), 42 comparisons (14 conditions × 3 genotypes) were made. For tethered LPC1-silencing walking experiments, 74 comparisons were made. For tethered LPC1-activation walking experiments, 20 comparisons were made. For imaging experiments, 12 comparisons each for LPC1 and LLPC1 were made. Statistical significance was noted with 1 star for p<0.05 and FDR q=0.05, and with 2 stars for p<0.01 and FDR q=0.01.

### Data and Code availability

We will make the data and code used to produce the major results of this study available at the time of publication. We will provide the most updated materials to correspond to the final version of the manuscript. FlyLight images and split-GAL4 driver lines will be available on the FlyLight website: https://splitgal4.janelia.org/cgi-bin/splitgal4.cgi. Data will be uploaded to FigShare, and analysis and plotting code will be available on https://github.com/reiserlab.

## Supplemental Information

Table S1. CSV file listing all of the gen 1 GAL4 lines (crossed to *UAS-Shibire*^ts1^) run through the Fly Vision Box screen, related to Fig. 1, S1. The table lists the summary data for each sequence of the Box protocol (see Methods for details). The first 729 entries are for the control genotype that was run throughout the GAL4 line screen.

**Movie S1: The *Drosophila* surgical preparation for 2-photon calcium imaging.**

